# Impact of astrocytic C3-production on neuronal mitochondrial dysfunction in tauopathy mouse models

**DOI:** 10.1101/2024.11.12.622892

**Authors:** Chenxu Lei, Bocheng Zhang, Junji Yamaguchi, Risako Tamura, Xingyu Cao, Yunhui Liu, Masahide Seki, Yutaka Suzuki, Kuninori Suzuki, Isei Tanida, Yasuo Uchiyama, Tatsuhiro Hisatsune

## Abstract

Neuronal mitochondrial dysfunction is associated with cognitive decline in neurodegenerative disorders such as Alzheimer’s Disease (AD). In this study, multiple pieces of evidence proved that phosphorylated tau (p-Tau) caused mitochondrial swelling and dysfunction in neurons. In a novel *in vitro* newborn neurons culture system, we discovered mitochondrial swelling and dysfunction were associated with increased p-Tau, leading to necroptosis activation, which was induced by Complement C3 (C3) produced from activated astrocytes. In the *in vivo* tauopathy mouse models, the effects of astrocytic C3 on tau-associated mitochondrial dysfunction and necroptosis were also discovered in hippocampal newborn neurons, and we directly showed that p-Tau aggregation was associated with mitochondria swelling in the hippocampal neurons by electron microscopy analysis. In addition, we proved the ability of compound anserine, which can block Tak1-Ikk dependent NF-κB activation, to further down-regulate astrocytic C3 production and alleviate neuronal mitochondrial dysfunction *in vitro* and *in vivo*, respectively. Down-regulation of astrocyte C3-production by anserine could also rescue mortality as well as cognitive and motor functions. Our findings first reported the contribution of p-Tau on neuronal mitochondrial dysfunction and proposed the therapies that down-regulate astrocytic C3 production have a potential role in alleviating this neurotoxic effect.

## Introduction

Alzheimer’s disease (AD), the most common form of dementia, is a neurodegenerative disease characterized by progressive neuronal loss and cognitive decline ^1^. The formation of neurofibrillary tangles (NFTs) composed of hyperphosphorylated tau (p-Tau) in the brain is one of the hallmarks of AD. Tau is a kind of microtubule (MT)-associated protein encoded by the *MAPT* gene, mainly expressed in neuronal axons, and has a primary function of promoting assembly and stability of microtubules ^2,3^. In tauopathies like AD, tau would become an abnormal hyperphosphorylated state, and is neurotoxic ^4,5^, linking to dysfunction of neuronal network and neurodegeneration ^6^. Clinical studies have reported that tau pathology is more associated with the cognitive decline and the progression of neurodegeneration in AD and other tauopathies compared with Aβ pathology ^7–9^. Recently, some molecules that are designed to target pathological tau have received positive results ^10,11^, indicating that therapies targeted tau-mediated neurodegeneration may be an ideal strategy for AD treatment.

Neuronal mitochondrial dysfunction is associated with neurodegeneration progression in tauopathies like AD ^12–14^. Evidence has shown that the aggregation of p-Tau in neurons would bring harmful effects on mitochondrial function ^15,16^. Neurons with pathological tau over-expression exhibited abnormal mitochondrial morphology, impaired mitochondrial dynamics, impaired oxidative phosphorylation and exacerbate oxidative damage ^17,18^. Besides, increased mitochondrial dysfunction is linked to neurodegeneration and neuronal death. Previous studies have report mitochondrial dysfunction would activate the necroptosis pathway by RIPK1/RIPK3/MLKL axis for the over- production of ROS ^19–21^, which has been reported to account for the death of neurons in AD ^22–24^. However, some detailed bases underlying the association between p-Tau, mitochondrial dysfunction and neurodegeneration remain unclear.

Astrocytes are the most abundant glial cells in the brain and play an important role in supporting neurons through astrocyte-neuron communications such as providing energy and metabolic supply ^25^. However, in tauopathies like AD, astrocytes would be over-activated with increased expression of diseased-associated genes and become neurotoxic ^26–28^. Activation of complement system is accompanied by astrocyte activation with over-production of complement C3, which is linked to exacerbated tau pathology through C3-C3aR signaling via GSK3β pathway ^29,30^. Astrocytic C3 over- production is associated with synapse loss, neuronal loss and cognitive decline in tauopathy models ^31,32^. Besides, C3 can negatively regulate the survival and migration of newborn neurons in adult mice ^33^. These studies suggest the harmful effects of astrocytic C3 on neuronal function and provide a possibility of the impact of C3 on neuronal mitochondrial dysfunction in tauopathies.

Here in this study, through using scanning electron microscope (SEM), we directly proved the contribution of p-Tau on neuronal mitochondrial swelling, both in the mature neurons and newborn neurons. We found that increased neuronal mitochondrial swelling would result in mitochondrial dysfunction and necroptosis activation, leading to cognitive decline and mortality. Besides, we found that down-regulate astrocytic C3 production by anserine (beta-alanyl-3-methyl-L-histidine), a natural anti-inflammatory imidazole dipeptide existing in vertebrate muscles that can target astrocytes ^34–36^, could alleviate tau-associated neuronal mitochondrial swelling. Down-regulation of astrocytic C3 production also rescued mortality as well as cognitive and motor functions in tauopathy mice. Our findings first revealed the impact of astrocytic C3 in mediating neuronal mitochondrial swelling, dysfunction and subsequent necroptosis activation in tauopathies and revealed the importance of therapies targeted astrocyte C3 production in alleviating this pathology.

## Results

### Pathological tau led to neuronal mitochondrial swelling in TE4 tauopathy model mice

To evaluate the role of pathological hyperphosphorylated tau on neuronal mitochondrial abnormity, we utilized human *APOE4* knock-in *MAPT* (P301S) transgenic mice (TE4). For the presence of *APOE4*, TE4 mice have been reported to show more severe tau-mediated neurodegeneration at about 9 months of age ^37^. However, there is lack of knowledge about the role of mitochondrial abnormity in neurodegeneration within this model. To directly observe the aggregation of p-Tau in neurons, we used correlative light and electron microscopy (CLEM). Through staining with phospho-Tau antibody AT8, which specifically recognizes tau phosphorylated at Ser202 and Thr205, we were able to observe the structure of AT8-positive neurons with aggregated tau fibers (Fig. 1a). Noticeably, we observed abnormal mitochondria with swollen structure in AT8-positive neurons, which were surrounded by tau fibers (Fig. 1a). For this, we compared the difference of mitochondrial structure in neurons with tau fibers aggregation and no tau fiber. As a result, although abnormal swollen mitochondria were commonly existed in hippocampus neurons in TE4 mice, we found a higher number of swollen mitochondria in neurons with tau fibers aggregation, which showed increased length and size (Fig. 1b and c). This result showed the contribution of p-Tau on neuronal mitochondrial swelling in TE4 tauopathy mice.

**Fig. 1:**
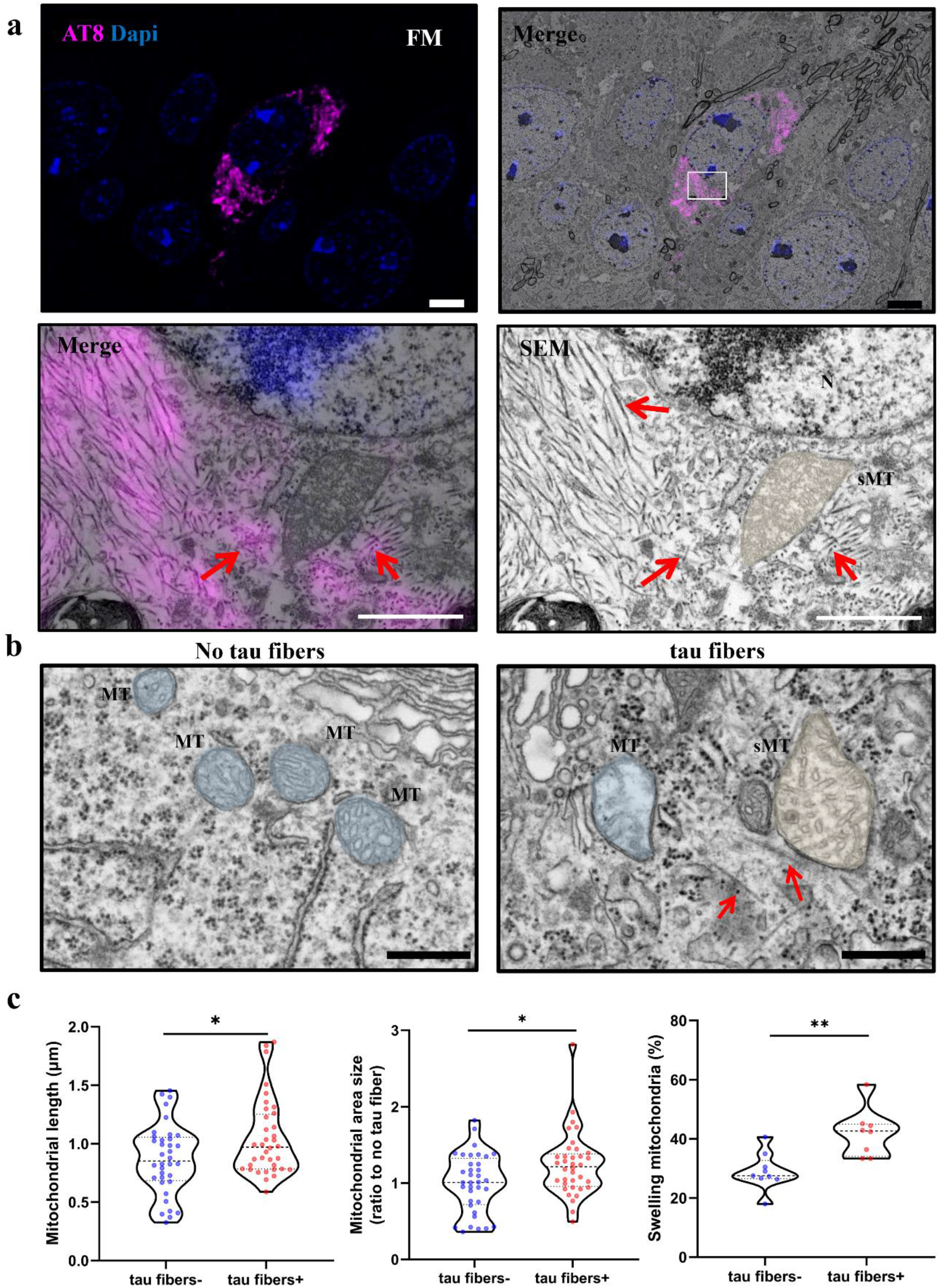
Neuronal mitochondrial swelling in TE4 (TauP301S *E4* knock-in) tauopathy model mice. **a** Representative CLEM (Correlative Light & Electron Microscopy) image of phosphorylated tau (AT8 antibody) positive neurons in the hippocampus of TE4 mice (scale bar: 1 μm). Dapi staining (blue) shows nucleus (N), and AT8 antibody staining (magenta) shows the area of p-Tau expression. Lower panels show enlarged images of upper panel (depicted by white rectangle). The red arrow in lower right panel shows aggregated-tau fibers in the cell body. Swelling mitochondria (sMT; >1 μm of long diameter; size volume increased; overlaid with yellow color), FM (Fluorescent Microscopy), SEM (Scanning Electron Microscopy). **b** Representative SEM image of neuron without tau fiber (left) and neuron containing tau fibers (right) in TE4 mice (red arrow: tau-fiber; scale bar: 0.5 μm). MT, normal mitochondria (blue), sMT, swelling mitochondria (yellow). **c** Quantification mitochondrial length of long diameter, mitochondrial area size and swelling mitochondria ratio of neurons with (8 neurons, 36 mitochondria) or without (9 neurons, 36 mitochondria) tau fibers in the hippocampus of TE4 mice. Data shows in the violin plots (median: bold dashed line; quarters (1/4 and 3/4: dashed lines), and were analyzed by Student’s t-test. \**p* < 0.05, \*\**p* < 0.01.

### Neuronal mitochondrial swelling was associated with neurodegeneration in TE4 mice

Neuronal mitochondrial abnormality is associated with neurodegeneration progression ^13,14^. In hippocampus area of TE4 mice, we observed degenerated neurons with abnormal structure, which contained swollen mitochondria and aggerated tau fibers (Fig. 2a). Interestingly, we observed necroptosis-like structure neurons in the hippocampus area of TE4 mice. These neurons showed obvious cell vacuolization and mitochondrial swelling (Fig. 2b). We also observed aggregated tau fibers in these neurons (Fig. 2b). This suggested the activation of neuronal necroptosis pathway in TE4 mice because of the effects of mitochondrial swelling caused by p-Tau. To further confirm this, we stained the tissues with necroptosis marker, p-MLKL. As a result, we observed p-MLKL-positive neurons and the co-location of p-Tau with p-MLKL in hippocampus area of TE4 mice (Fig. 2c and d). P-Tau is regarded as a main trigger for neuronal necroptosis ^21,38^, our data confirmed this and indicated it was associated with neuronal mitochondrial swelling.

**Fig. 2:**
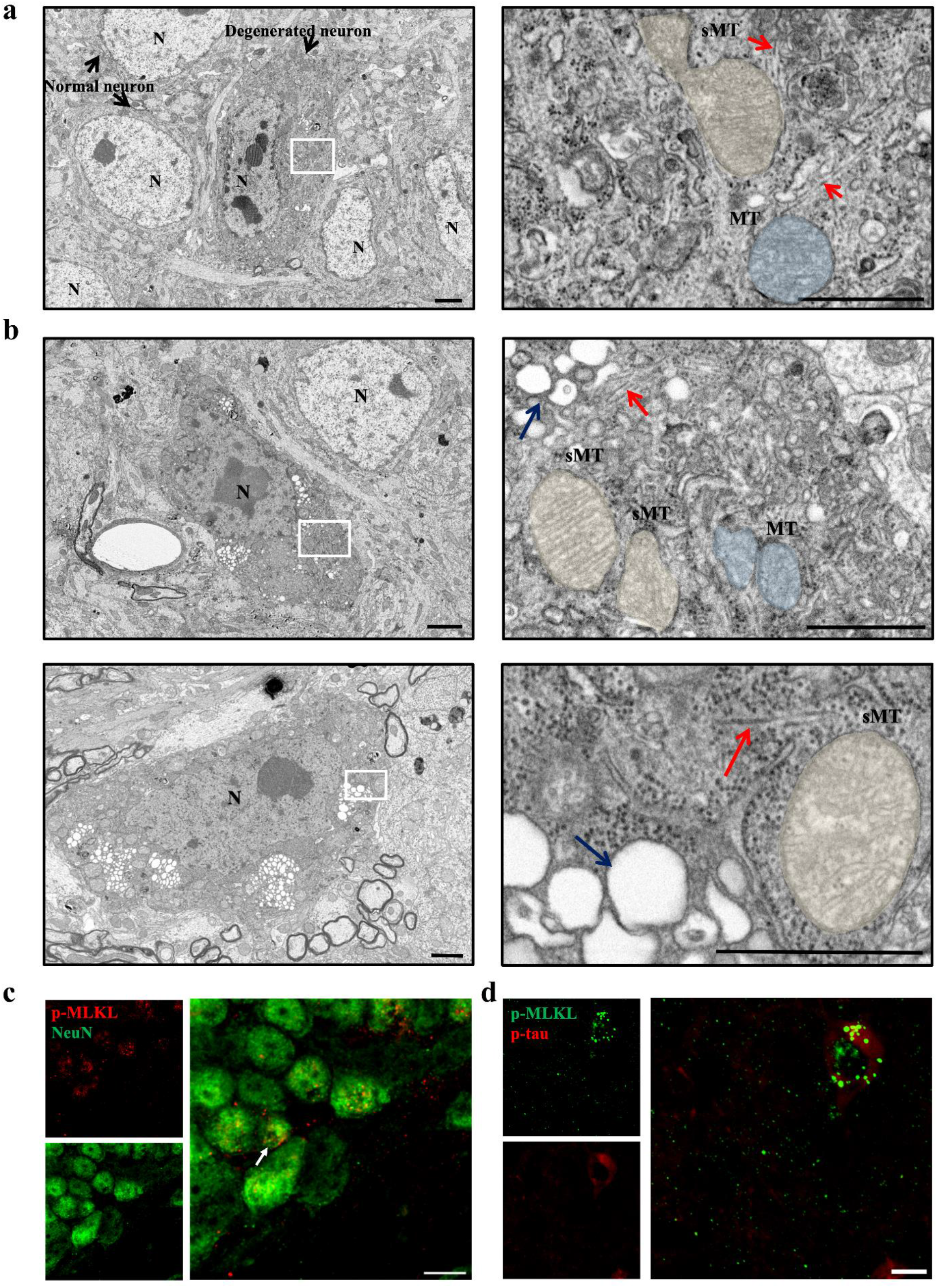
Neuronal mitochondrial swelling exhibiting necroptosis-like structures in TE4 mice. **a** Representative SEM images of hippocampal neurons in TE4 mice (Scale bar: 1 μm). Nucleus of normal neurons are bright, but nucleus of degenerated neurons become dark. N, nucleus, MT, normal size mitochondria (blue), sMT, swelling mitochondria (yellow). The red arrow showed tau fibers. **b** Representative SEM image of hippocampal neurons in TE4 mice with necroptosis-like phenotypes of dark nucleus and robust cellular vacuolization, (Scale bar: 1 μm). The red arrow showed tau fibers, and blue arrow showed the cell vacuolization. **c** Representative images of p-MLKL staining neurons (NeuN) in the hippocampus of TE4 mice, white arrow showed the p-MLKL-positive neuron (scale bar: 10 μm). **d** Representative necroptosis-like hippocampal neuron with of p-MLKL pyramidal cytoplasmic staining colabelled with p-Tau (AT8) in TE4 mice (scale bar: 10 μm).

### Down-regulation of astrocytic C3 production alleviated neuronal mitochondrial swelling in TE4 mice

Recent years, complement system over-activation has been reported to promote tau pathology and neurodegeneration through C3-C3aR signaling ^29–31^. In our previous study, we found a natural anti-inflammatory compound named anserine, had the ability to suppress astrocyte activation and protect cognitive function ^34,39^. Therefore, we thought anserine may have the ability to down-regulate astrocytic C3 production and alleviate tau-associated neuronal mitochondrial swelling. For this, we established a novel cell culture model using an immortalized neural stem cell line called MSP-1 cells. MSP-1 cells lack *P53* and were allowed to proliferate indefinitely without apoptosis and were allowed to be induced to differentiate into neurons and astrocytes as we previously described ^40–44^. We induced MSP-1 cells to differentiate into astrocytes by LIF and CNTF, and to simulate the inflammatory environment, we activated the MSP-1 astrocytes with TNF-α (Fig. 3a). TNF-α treatment significantly activated the MSP-1 astrocytes by up-regulating NF-κB signaling pathway (Fig. 3b and c). At the same time, MSP-1 astrocytes pre-treated with anserine showed decreased NF-κB activation (Fig. 3b and c). Through studying the changes of NF-κB upstream signals, the results showed that anserine could suppress Tak1 phosphorylation to prevent NF-κB activation by Tak1-Ikk axis (Fig. 3g-l). For the ability to suppress NF-κB activation, anserine treatment significantly reduced the expression of pro-inflammatory factors represented by *C3* (Fig. 3d-f). These results suggested the ability of anserine on down-regulating astrocyte activation and C3 production.

**Fig. 3:**
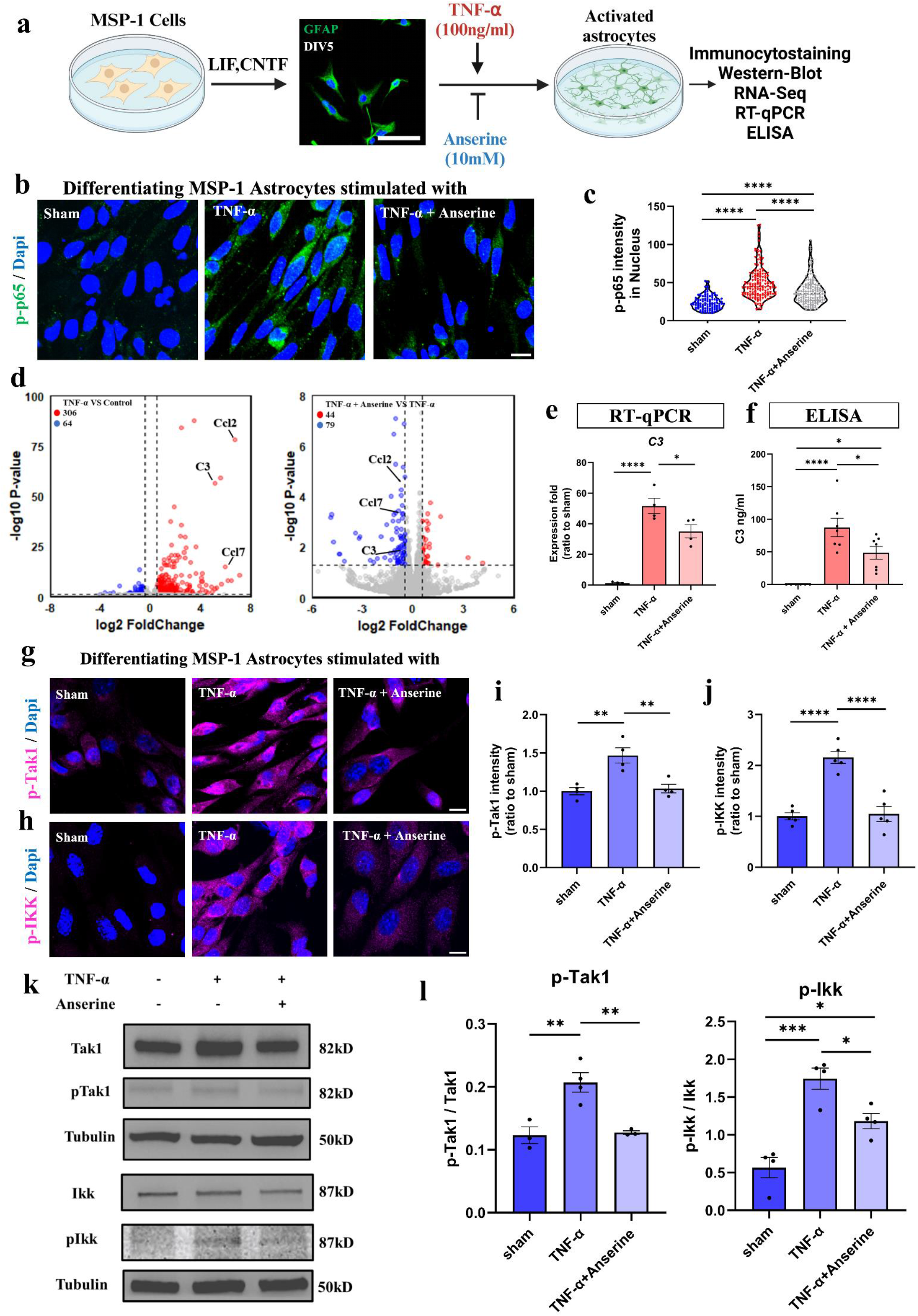
Down-regulation of astrocytic C3 production by anti-inflammatory peptide, anserine via blocking Tak1-Ikk dependent NF-κB activation. **a** Brief description of experiment confirming down-regulation of astrocytic C3 production by a natural anti-inflammatory peptide, anserine. MSP-1 astrocytes were differentiated from neural stem cell (MSP-1) through the stimulation of CNTF and LIF for 5 days *in vitro* (scale bar: 50 μm). TNF-α stimulation was used to induce astrocyte activation and C3 production in MSP-1 astrocytes in 24-hour culture. To study the effects of anserine on NF-κB activation, TNF-α treatment was performed for 30 min. **b** Representative images of p-p65 staining in MSP-1 astrocytes of sham, TNF-α and Anserine + TNF-α group (scale bar: 10 μm). **c** Quantification of p-p65 intensity in the nucleus of 3 groups, (*n*=110- 152). **d** Volcano plot of identified DEGs (|log2 Fold-change|> 0.5, p < 0.05, up, red, down, blue) between TNF-α group and Control group (left) or Anserine + TNF-α group and TNF-α group (right). **e** Quantification of *C3* expression of 3 groups, (*n*=3-6). **f** Result of C3 concentration in cultured medium of sham, TNF-α and Anserine + TNF-α group, (*n*=6-7). **g** and **h** Representative image of p- Tak1 (**g**) and p-Ikk (**h**) staining in MSP-1 astrocytes (scale bar: 10 μm). **i** and **j** Quantification of p- Tak1 (**i**) and p-Ikk (**j**) intensity in each group, (*n*=4-5). **k** Representative western blot images of Tak1, p-Tak1, IKK and p-IKK in each group. **l** Quantification of relative expression in each group, (*n*=3-4). Data represent means ± SEM and were analyzed by one-way ANOVA with Tukey’s multiple comparisons test. \**p* < 0.05, \*\**p* < 0.01, \*\*\**p* < 0.001, \*\*\*\**p*<0.0001. Some elements of figure (**a**) were created in https://BioRender.com.

Next, we treated TE4 mice with anserine from 7 months of age for 8 weeks, called TE4 (A) mice. Consistent with the *in vitro* study, through RNA-Sequencing (RNA-Seq) analysis using MACS-isolated astrocytes, we found that anserine treatment down-regulated the expression of A1-specific astrocyte markers represented by *C3* (Fig. 4a and b and Supplementary Fig. 1a-d). Besides, *Mfeg8* that down-regulated A1 astrocyte activation and C3 production was significantly down-regulated in astrocytes of TE4 mice and up-regulated after anserine treatment ^45^ (Fig. 4a and b). Down-regulation of C3 production in TE4 mice was confirmed by RT-qPCR, ELISA and immunostaining (Fig. 4c-f). Further analysis of common DEGs that up-regulated in TE4 group and down-regulated in TE4 (A) group by Go Term and KEGG analysis revealed that anserine treatment significantly inhibited the complement activation by NF-κB pathway (Supplementary Fig. 1e and f). Then we studied the effects of astrocytic C3 down-regulation on neuronal mitochondrial swelling in TE4 mice by scanning electron microscope (SEM). As a result, we found a reduced number of abnormal swollen mitochondria with reduced mitochondrial area and length in hippocampus neurons in TE4 (A) mice (Fig. 4g and h and Supplementary Fig. 2). Through AT8 staining and PT181 (recognize tau phosphorylated at Thr181) staining, it was shown that TE4 (A) mice exhibited attenuated tau pathology both in CA1 and DG area (Supplementary Fig. 3a-d). Noticeably, the expression of C3 in GFAP-positive cells had a strong association with tau pathology in TE4 mice (Supplementary Fig. 3e and f), which suggested the contribution of astrocytic C3 on promoting tau pathology and neuronal mitochondrial swelling. Besides, because of alleviated mitochondrial swelling, TE4 (A) mice showed attenuated neurodegeneration that was shown as increased neuronal density (Supplementary Fig. 3e- g), which may a consequence of reduced necroptosis activation caused by mitochondrial swelling.

**Fig. 4:**
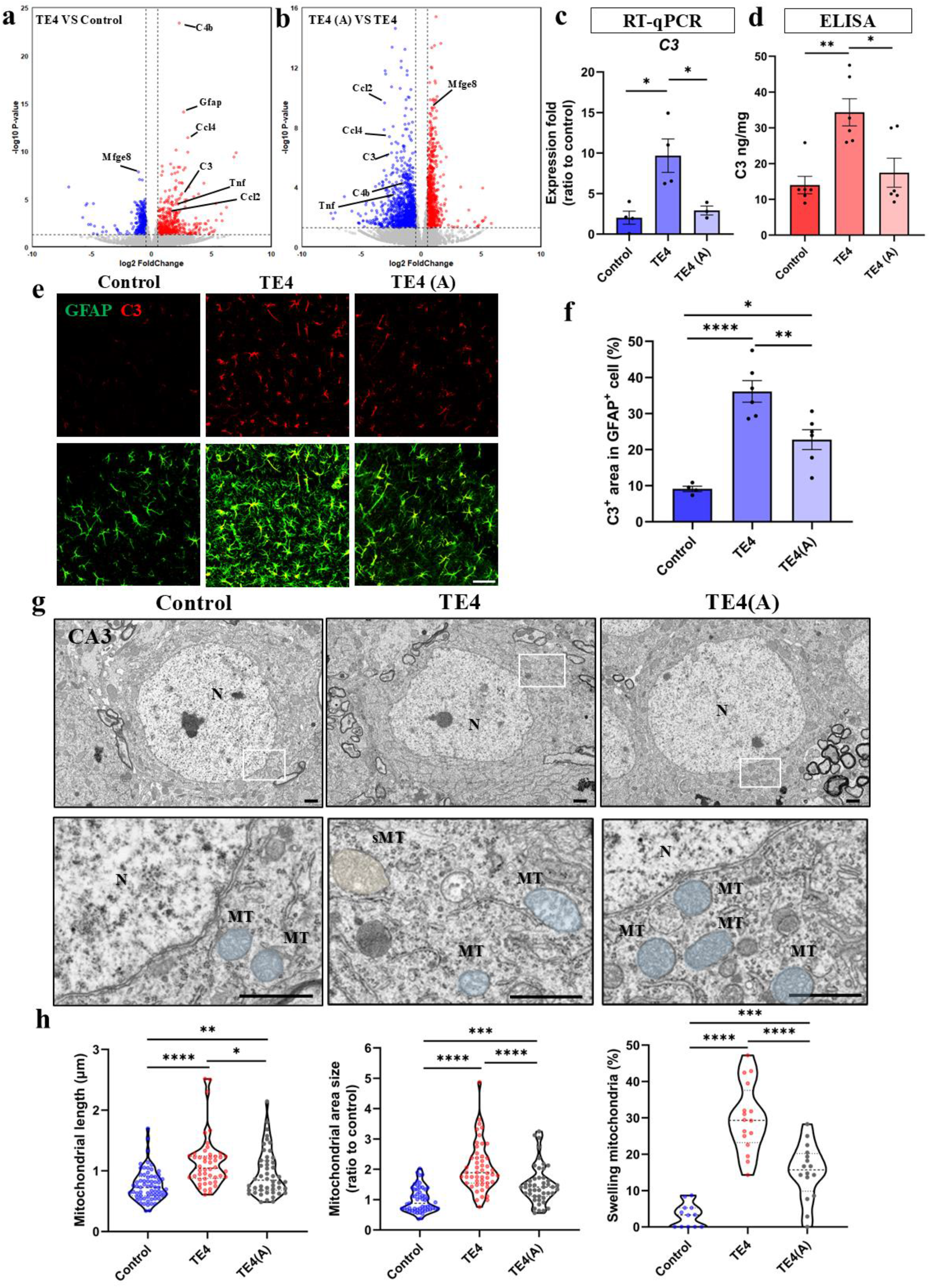
Down-regulation of astrocytic C3 production alleviated mitochondrial swelling in TE4 mice. **a** The description of astrocytes isolation for RNA-Seq used ACSA-2 Microbead from mouse brain. **a** and **b** Volcano plot of identified DEGs (|log2 Foldchange|>1, *p*<0.05, up, red, down, blue) between astrocytes of TE4 mice and Control mice (**b)** or between astrocytes of TE4 (A) mice (TE4 mice treated with anserine) and TE4 mice (**b**) (see legend of Fig. 5a). **c** RT-qPCR results of the expression of *C3* in each group (*n*=3-4). **d** Results of C3 concentration in hippocampus of 3 groups of mice, (*n*=6). **e** Representative images of GFAP and C3 co-staining in the hippocampus area of 3 groups of mice (scale bar: 50 μm). **f** Quantification of C3-positive area in GFAP-positive cells in each group, (*n*=4-6). **g** Representative SEM images of hippocampal neurons in control, TE4 and TE4 (A) mice (scale bar: 1 μm). N, nuclear, MT, normal mitochondria (blue), sMT, swollen mitochondria (yellow). **h** Quantification mitochondrial length, mitochondrial area size and swelling mitochondria ratio in CA3 area of 3 groups of mice (neuron number: 12-16, mitochondria number: 46-67). Data represent means ± SEM (**e, d and f**) or showed in the violin plots (median: bold dashed line; quarters (1/4 and 3/4: dashed lines) (**h**) and were analyzed by one-way ANOVA with Tukey’s multiple comparisons test. \**p* < 0.05, \*\**p* < 0.01, \*\*\**p* < 0.001, \*\*\*\**p* < 0.0001.

### Suppressed astrocytic C3 production reduced mortality and improved cognitive function in TE4 mice

P301S tau model mice would exhibit paralysis associated with a hunched posture during tau pathology progression, which results in feeding inability and death ^6^. Since anti-inflammatory therapy prolonged the survival rate in P301S mice ^6^, we compared the survival rate between TE4 mice and TE4 (A) mice from the beginning of anserine treatment and we observed that TE4 (A) mice hardly developed a hunched posture and could easily live over 10 months of age, while only about 55% TE4 mice could alive over 10 months of age (Fig. 5a-c and supplementary video. 1). Anserine treatment also prevented the weight decline in TE4 mice (Fig. 5d). For the paralysis, TE4 mice also showed impairment of motor functions such as grip strength decline. Compared with TE4 mice, the grip strength of TE4 (A) mice was consistent with control mice (Fig. 5e).

**Fig. 5:**
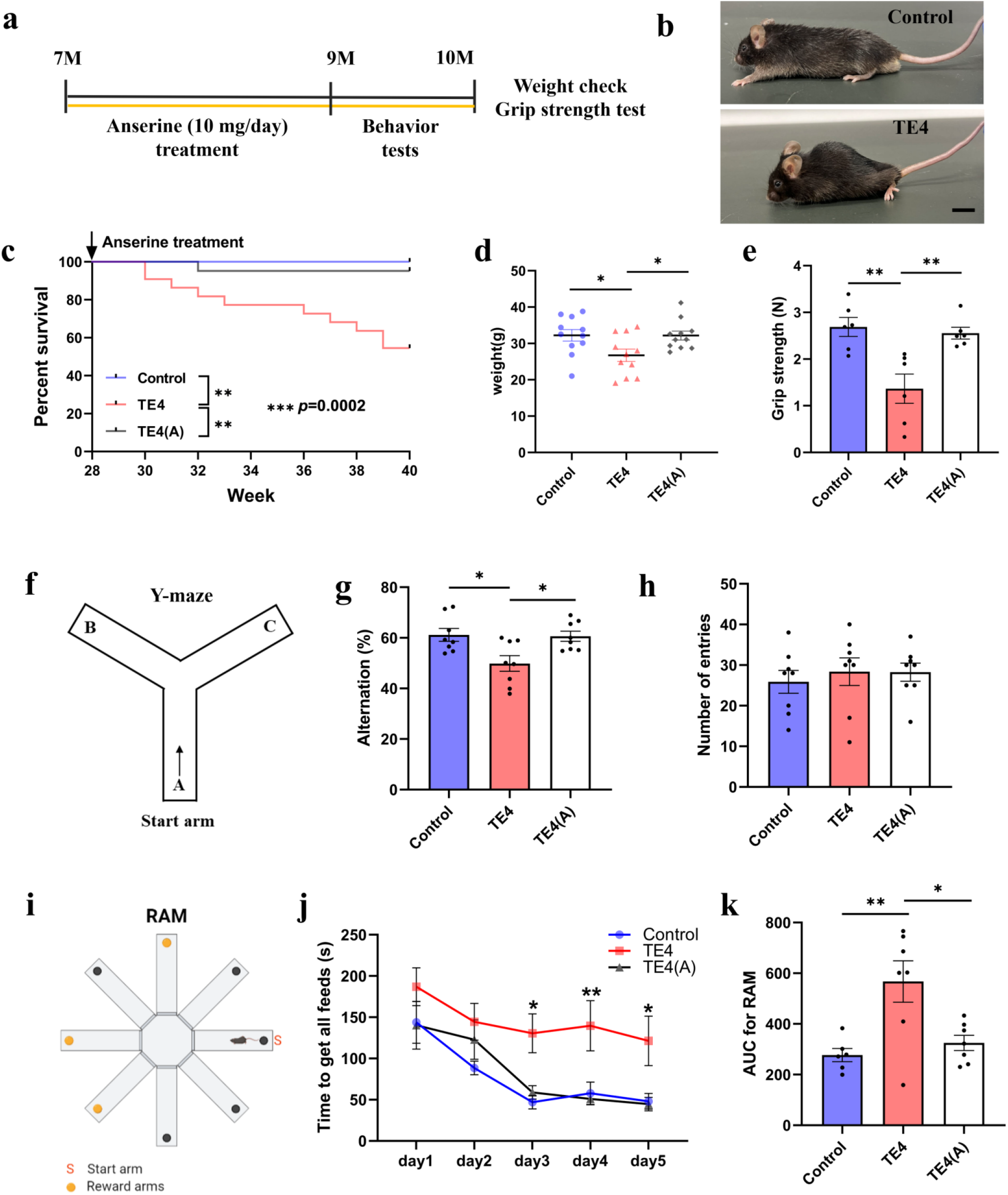
Impact of astrocytic C3 production on mortality and cognitive functions of TE4 mice. **a** Description of behavior experiments and physiological function check. **b** Representative images of control mice and TE4 mice (scale bar: 1 cm). **c** Kaplan-Meier survival curve of control, TE4 and TE4 (A) groups, (*n*=15-22). Kaplan-Meier survival analysis was performed by log-rank (Mantel-Cox) test, *p*=0.0002. Control and TE4, *p*=0.0031; TE4 and TE4 (A), *p*=0.0027. **d** Quantification of body weight of 3 groups of mice at the end of behavior tests, (*n*=11). **e** Quantification of grip strength of 3 groups of mice at the end of behavior tests, (*n*=6). **f** Description of Y-Maze, the mouse would start at A site. **g** and **h** Results of Y-maze. The results were presented as alternation rates (%) (**g**) and the number of entries (**h**) in the Y-maze of 3 groups of mice during 10 min, (*n*=8). **i** Description of Radial arm maze (RAM). **j** and **k** Results of RAM. The time of cost to find all the feeds was recorded (**j**) in a 5-days RAM test. At the same time, the area of the time curve for the 5-day RAM test was calculated for each mouse (**k**) (*n*=6-7). Data represent means ± SEM and were analyzed by one-way ANOVA with Tukey’s multiple comparisons test (**d**, **e**, **g**, **h**, and **k**) or two-way repeated ANOVA with Tukey’s multiple comparisons test (**j**). \**p* < 0.05, \*\**p* < 0.01. Some elements of figure (**i**) were created in https://BioRender.com.

Then we evaluated the cognitive function in TE4 mice after anserine treatment through Y-maze and Radial arm maze (RAM) tests. In Y-maze, TE4 mice showed a significant decrease in alternation rate compared with control mice. Besides, TE4 mice take more time to find all the rewards in the RAM test (Fig. 5f-k). These results indicated that neuronal necroptosis led to a decline of cognitive function in TE4 mice. Compared with TE4 mice, TE4 (A) mice showed an increased alternation rate in Y-maze (Fig. 5g), and no difference was found in the total number of entries (Fig. 5h). Similar results were found in the RAM test that TE4 (A) mice showed a shorter time to find all the rewards compared with TE4 mice (Fig. 5i-k). These findings indicated that suppressed astrocyte activation by anserine rescued the cognitive function and kept TE4 mice in a healthy state, which may be associated with attenuated tau pathology because of the down-regulated C3 production.

### Loss of function of down-regulating astrocytic C3 production caused tau pathology and neurodegeneration

To further confirm the effect of astrocytic C3 on tau-associated neurodegeneration, we knock down (KD) the *Pept2*, a transporter of anserine that only expresses in astrocytes by AAV-shPEPT2- PHP.eB ^46^ (Fig. 6a-c). We expected anserine can be taken by astrocytes and directly affect astrocytes. Interestingly, compared with TE4 (A) mice injected with AAV-Scramble RNA-PHP.eB (TE4 (A)- shCon), TE4 (A) mice with AAV-shPEPT2-PHP.eB injection (TE4 (A)-shPEPT2) exhibited an increased astrocyte activation and C3 expression in GFAP-positive astrocytes (Fig. 6d-f). These results suggested that *Pept2* KD blocked the effects of anserine on A1 astrocyte activation. As a result, TE4 (A)-shPEPT2 mice showed increased p-Tau expression and increased number of p-MLKL positive neurons compared with TE4 (A)-shCon mice (Fig. 6g-j). We also investigated the cognitive function between TE4 (A)-shPEPT2 mice and TE4 (A)-shCon mice. Noticeably, TE4 (A)-shPEPT2 showed the same alternation rate as TE4 mice in the Y-maze test, while TE4 (A)-shCon mice also showed improved cognitive function (Fig. 6k). These findings confirmed that astrocytic C3 production may be the main reason of tau-associated neurodegeneration and necroptosis activation. Anserine, which can target astrocytes by *Pept2*, has the ability to down-regulate astrocytic C3 production to prevent the neurotoxic effects (Fig. 6l).

**Fig. 6:**
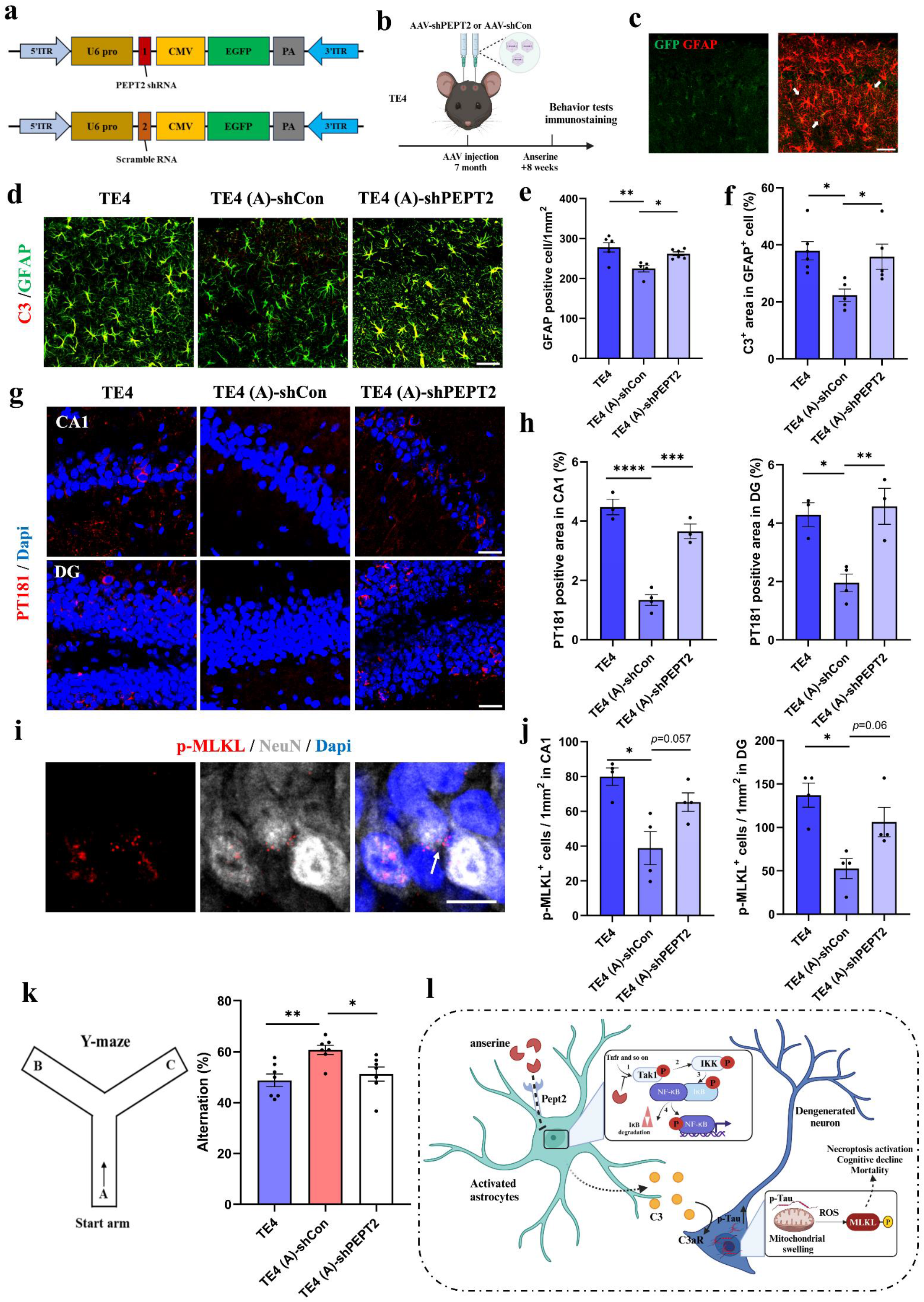
Blocking of astrocytic C3 production down-regulation caused neurodegeneration in TE4 mice. **a** and **b** The description of blocking of astrocytic C3 production down-regulation by an *in vivo* shRNA experiment. To block astrocytic intracellular mobilization of anserine, shRNA for specific transporter protein, PEPT2, was constructed in AAV (adeno-associated viral)-vector and inoculated into TE4 mice. **c** Representative images of astrocytic expression of PEPT2-shRNA-AAV-vector, in which vector-driven GFP was expressed in GFAP-positive astrocytes in the hippocampus area of AAV-injected TE4 mice (scale bar: 50 μm). **d** Representative images of GFAP and C3 co-staining in hippocampus area in TE4, TE4 (A)-shCon and TE4 (A)-shPept2 mice (scale bar: 50 μm). **e** Quantification of the number of GFAP positive-cells in each group, (*n*=5-6). **f** Quantification of C3-positive area in GFAP-positive cells in each group, (*n*=5-6). **g** Representative images of PT181 staining in hippocampal CA1 and DG area in TE4, TE4 (A)-shCon and TE4 (A)-shPept2 mice (scale bar: 25 μm). **h** Quantification of PT181- positive stain area in each group, (*n*=3-4). **i** Representative images of p-MLKL and NeuN co-staining in TE4 mice (scale bar: 10 μm), white arrow showed the p-MLKL-positive neuron. **j** Quantification of p-MLKL-positive neurons in CA1 and DG area in each group, (*n*=4). **k** Result of Y-maze in TE4, TE4 (A)-shCon and TE4 (A)-shPept2 mice, (*n*=7). **l** Brief diagram about the mechanism of anserine suppressing astrocytic C3 to alleviate neuronal mitochondrial swelling. Data represent means ± SEM and were analyzed by one-way ANOVA with Tukey’s multiple comparisons test. \**p* < 0.05, \*\**p* < 0.01, \*\*\**p* < 0.001, \*\*\*\**p* < 0.0001. Some elements of figure (**b and l**) were created in https://BioRender.com.

### Down-regulation of astrocytic C3 production alleviated tau-associated neuronal mitochondrial swelling and dysfunction *in vitro*

Subsequently, to further confirm this, we induced MSP-1 cells to differentiate into neurons and treated the MSP-1 neurons with the culture medium of MSP-1 astrocytes induced by TNF-α (Fig. 7a). In DIV5, almost all the MSP-1 cells expressed Map-2 and also expressed immature neuron marker PSA-NCAM (Supplementary Fig. 4). Therefore, these cells can regard as MSP-1 newborn neurons. As a result, through staining with MitoBright IM and using SEM, we found that mitochondria in MSP- 1 neurons treated with culture media (CM) from TNF-α group (TNF CM) showed swollen structure with increased length and size (Fig. 7b-e). We further found that increased mitochondrial swelling led to decreased mitochondrial membrane potential and increased ROS production (Fig. 7f-i), indicating the mitochondrial swelling in MSP-1 neurons was accompanied by impaired mitochondrial function. TNF CM treatment also increased p-Tau expression that was shown in AT8 staining (Supplementary Fig. 5c and d). Compared with TNF CM treated group, we found that MSP-1 neurons treated with CM from the TNF-α+Anserine group (TNF +Anserine CM) showed reduced number of swollen mitochondria and alleviated mitochondrial dysfunction (Fig. 7b-i). Noticeably, mitochondrial swelling and dysfunction caused by TNF CM was also significantly prevented by C3aR antagonist (Fig. 7b-i). Besides, MSP-1 neurons treated TNF +Anserine CM or TNF CM-C3aRA showed significant decreased AT8 intensity (Supplementary Fig. 5c and d). Consistent with our findings in TE4 mice, these results indicated the crucial role of C3 to induce tau-associated mitochondrial swelling in MSP- 1 neurons.

**Fig. 7:**
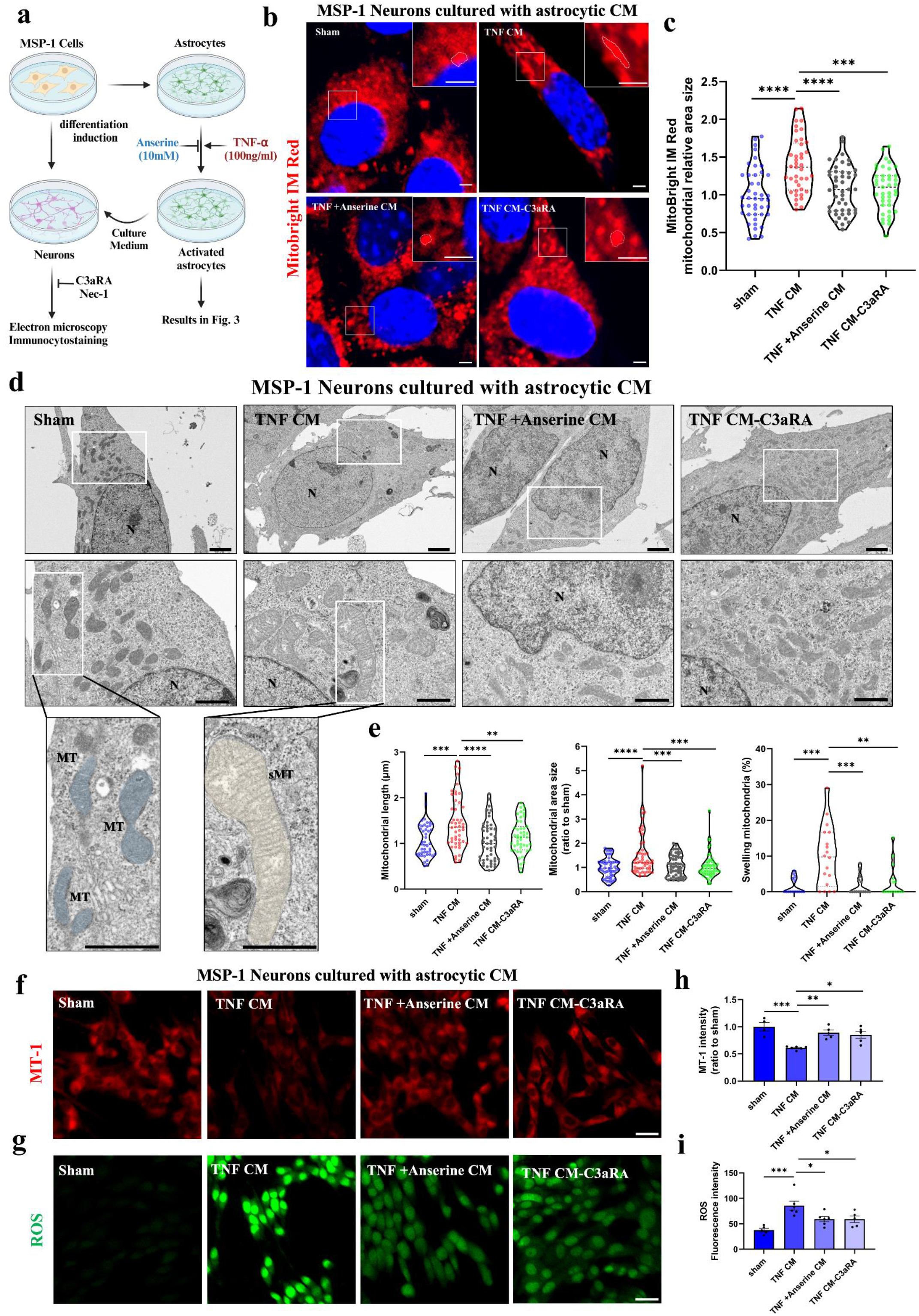
Astrocytic C3 induced mitochondrial swelling and dysfunction in MSP-1 newborn neurons. **a** Brief description of *in vitro* study using differentiating MSP-1 newborn neurons. MSP-1 neurons were treated with collected Cultured medium (CM) from MSP-1 astrocytes for 24 hours. **b** Representative images of MSP-1 neurons stained with Mitobright IM Red (scale bar: 2 μm). **c** Quantification of mitochondrial area size of each group (*n*=41-47). **d** Representative electron microscopic images of MSP-1 neurons treated with sham, TNF CM, TNF +Anserine CM and TNF CM-C3aRA (scale bar: 2 μm (up), 1 μm (down)). N, nuclear, MT, normal mitochondria (blue), sMT, swollen mitochondria (yellow). **e** Quantification mitochondrial length, mitochondrial area size and swelling mitochondria ratio in each group (neuron number: 15-18, mitochondria number: 47-53). **f** Representative images of MT-1 staining of MSP-1 neurons treated with sham, TNF CM, TNF +Anserine CM and TNF CM-C3aRA (10μM) (scale bar: 25 μm). **g** Quantification of MT-1 intensity in each group, (*n*=4-6). **h** Representative images of ROS staining of MSP-1 neurons treated with sham, TNF CM, TNF +Anserine CM and TNF CM-C3aRA (10μM) (scale bar: 25 μm). **i** Quantification of ROS fluorescence intensity in each group, (*n*=5-6). Data represent means ± SEM (**h and i**) or showed in the violin plots (median: bold dashed line; quarters (1/4 and 3/4: dashed lines) (**c and e**) and were analyzed by one-way ANOVA with Tukey’s multiple comparisons test. \**p*<0.05, \*\**p* < 0.01, \*\*\**p* < 0.001. \*\*\*\**p*<0.0001. Some elements of figure (**a**) were created in https://BioRender.com.

We also investigated the impact of astrocytic C3 on neurodegeneration in MSP-1 neurons. Consistent with our results in TE4 mice, we observed an increased p-MLKL intensity in MSP-1 neurons treated with TNF CM (Supplementary Fig. 5a and b), while MSP-1 neurons treated with TNF +Anserine CM or TNF CM-C3aRA showed reduced p-MLKL intensity to the same level as the MSP- 1 neurons treated with Necrostatin-1 (Nec-1), a necroptosis inhibitor (Supplementary Fig. 5a and b), which revealed that C3 was the crucial factor in necroptosis activation. To further confirmed this, we treated MSP-1 neurons with C3. C3 treatment also increased AT8 intensity in MSP-1 neurons, which led to decreased mitochondrial membrane potential and increased p-MLKL intensity (Supplementary Fig. 5e-i). Noticeably, MSP-1 neurons treated with C3 showed similar p-MLKL intensity to MSP-1 neurons treated with TNF CM (Supplementary Fig. 5c and d). Taken together, these results suggested the impact of astrocytic C3 production on tau-associated neuronal mitochondrial swelling, which would cause mitochondrial dysfunction and necroptosis activation.

### Down-regulation of astrocyte C3 production alleviated mitochondrial dysfunction in newborn neurons in E4-3Tg model mice

During AD progression, neuronal dysfunction and degeneration is also occurred in the newborn neurons ^47^. Recently, Wang and colleagues reported that the activation of astrocytes may affect the number of immature neurons in the aged human hippocampus and they found some key mediators like NF-κB and STAT3 signaling that may inhibit adult hippocampal neurogenesis (AHN) ^48^. Besides, C3 has been reported to affect the survival of newborn neurons ^33^. Since MSP-1 neurons can be regard as an *in vitro* newborn neurons model for the expression of PSA-NCAM (Supplementary Fig. 4), we supposed that astrocytic C3 over-production had the same effects on newborn neurons *in vivo*. Therefore, finally, we studied the effects of astrocyte activation and C3 production on newborn neurons. Since tauopathy mice were mostly used to study the neurodegeneration, and there remained contention about the role of tau pathology in neurogenesis ^49^, we utilized *APOE4* knock-in/APP/PS1/P301S transgenic (E4-3Tg) model mice in which the number of newborn neurons was significantly decreased at 6 months of age (Fig. 8h-k). To clarify the mechanism, we isolated newborn neurons by MACS using anti-PSA-NCAM microbeads from E4-3Tg mice at 6 months of age and did RNA-Seq analysis (Fig. 8a). As a result, we noticed that expression of genes related to mitochondria, oxidative phosphorylation and electron transport chain was down-regulated and genes related to oxidative damage response was up-regulated in newborn neurons of E4-3Tg mice (Fig. 8b and c). These results revealed that mitochondrial dysfunction was also occurred in the newborn neurons in E4-3Tg mice. Besides, genes related to synapse plasticity and neuron maturation were also down- regulated, represented by *Syn, shank2*, *Nrn1, NeuronD6* and *Map1a* (Supplementary Fig. 6). This indicated that mitochondrial dysfunction led to the impairment of maturation process as energy supple is important in this process. In addition, newborn neurons in anserine-treated E4-3Tg mice, E4-3Tg (A) mice, did not show severe mitochondria dysfunction and increased oxidative damage (Fig. 8b and c). Anserine treatment also protected the synapse plasticity and neuronal maturation in newborn neurons (Supplementary Fig. 6), which may relate to the decreased p-Tau expression in DCX-positive cells because of the down-regulation of astrocytic C3 production (Fig. 8d and e and Supplementary Fig. 7). As a result, we found an increased number of newborn neurons in E4-3Tg (A) mice (Fig. 8h-k). These findings indicated the dysfunction of newborn neurons such as impaired synapse plasticity and mitochondrial dysfunction mediated by tau pathology in E4-3Tg mice and the ability of anserine to alleviate this through down-regulating C3 production.

**Fig. 8:**
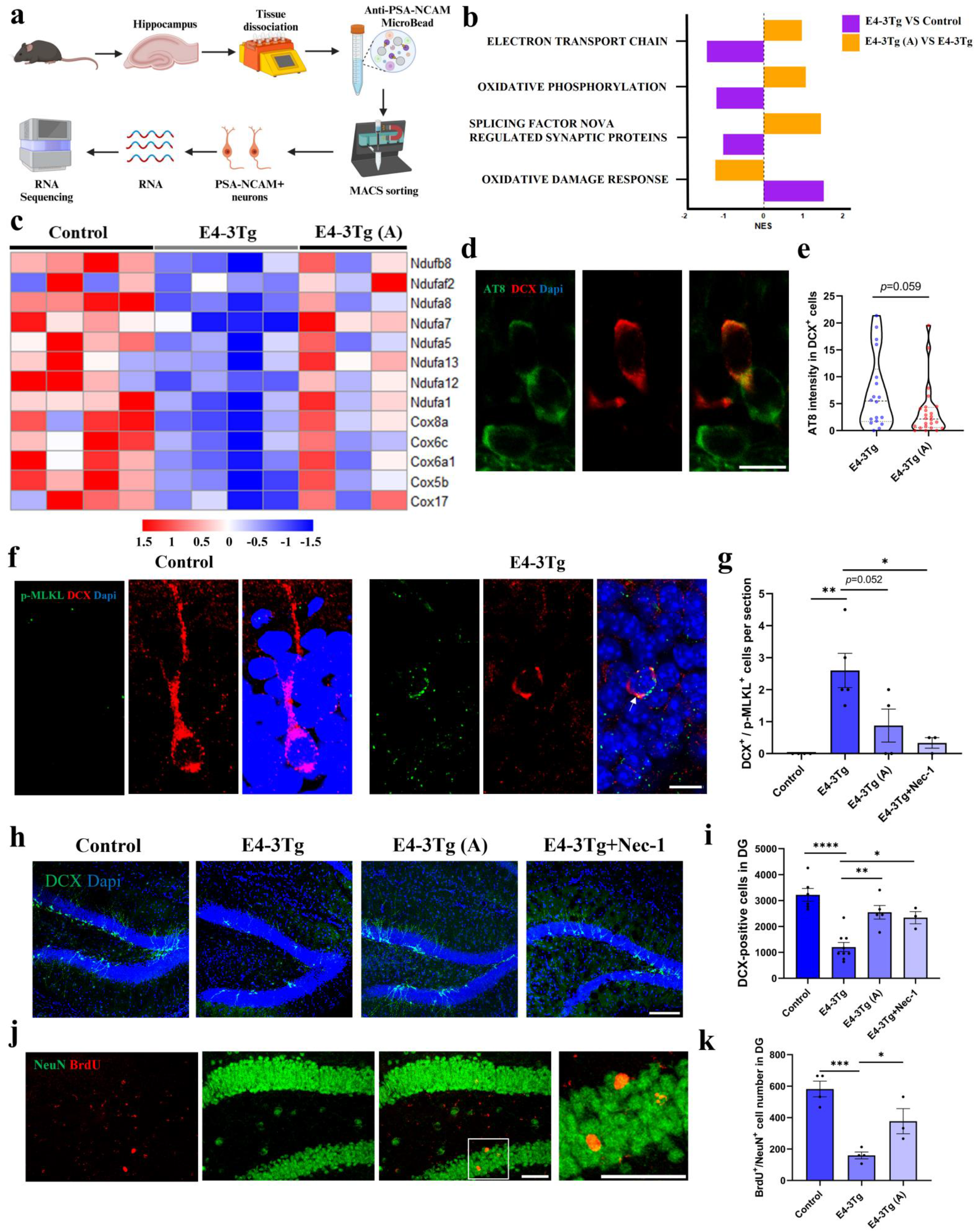
Down-regulation of astrocytic C3 production alleviated mitochondrial dysfunction in newborn neurons in E4-3Tg model mice. **a** Brief description of newborn neuron isolation for RNA-Seq analysis using PSA-NCAM Microbead from mouse hippocampus. **b** Representative GESA results using the Wiki Pathway database focused on mitochondrial function. Purple, E4-3Tg VS control. Orange, E4-3Tg (A) VS E4-3Tg. **c** Heatmap that shows expression levels of mitochondria markers of 3 groups of mice. All the selected genes were significantly down-regulated in E4-3Tg group and up-regulated in E4-3Tg (A) group. **d** Representative images of DCX and AT8 co-staining in DG area of E4-3Tg mice and E4-3Tg (A) mice (scale bar: 10 μm). **e** Quantification of AT8 intensity in DCX positive cells of each group, (*n*=18-24). **f** Representative image of DCX and p-MLKL co-staining in DG area of control mice and E4-3Tg mice (scale bar: 10 μm). **g** Quantification of the number of DCX^+^/ p-MLKL^+^ cells in each group (*n*=3-5). **h** Representative image of DCX staining in DG area of control mice, E4-3Tg mice, E4-3Tg (A) mice and Nec-1 treated E4-3Tg mice (scale bar: 100 μm). **i** Quantification of DCX-positive cell number in DG in each group, (*n*=3-8). **j** Representative image of BrdU and NeuN co-staining in the DG area (scale bar: 50 μm). **k** Quantification of BrdU^+^/NeuN^+^ cell number in DG of each group, (*n*=3-4). Data represent means ± SEM (**g, i** and **k**) or showed in the violin plots (median: bold dashed line; quarters (1/4 and 3/4: dashed lines) (**e**) and were analyzed by Student’s *t*-test (**e**) and one-way ANOVA with Tukey’s multiple comparisons test (**g, i** and **k**). \**p* < 0.05, \*\**p* < 0.01, \*\*\**p*<0.001. Some elements of figure (**a**) were created in https://BioRender.com.

We also studied whether mitochondrial dysfunction would cause necroptosis activation in newborn neurons. Noticeably, through immunostaining, we observed the expression of p-MLKL in DCX-positive cells, which can be prevented by Nec-1 treatment (Fig. 8f and g). Nec-1 treatment also significantly increased the number of DCX-positive cells (Fig. 8h and i). These results revealed the activation of the necroptosis pathway in newborn neurons of E4-3Tg mice. We also found the reduced number of p-MLKL^+^/DCX^+^ cells after anserine treatment (Fig. 8f and g), which suggested that the suppression of astrocyte activation and C3 production was linked to improved AHN levels and reduced loss of newborn neurons caused by necroptosis activation. Taken together, our findings showed that astrocyte activation and C3 over-production were also related to mitochondrial dysfunction and necroptosis activation in newborn neurons.

### Suppressed astrocyte activation reduced mortality and improved cognitive function in E4-3Tg mice

Because of the tau pathology, E4-3Tg mice also a showed reduced survival rate (Supplementary Fig. 8a). We also investigated whether the anserine treatment had the same effects to prolong the survival rate of E4-3Tg mice just like what we found in TE4 mice. Our data showed that anserine also prevented the mortality of E4-3Tg mice (Supplementary Fig. 8a). Besides this, we found improved cognitive function in E4-3Tg (A) mice in Y-maze and CFC test (Supplementary Fig. 8b-e). Consistent with TE4 mice, these results suggested that improved AHN levels by down-regulating astrocytic C3 production rescued the cognitive function as well as prolonged the survival rate in E4-3Tg mice.

## Discussion

Neuronal mitochondrial abnormality and dysfunction is one of the hallmarks of dementia like AD ^13,14^. As highly energetic cells, neurons need healthy mitochondria to maintain the neuronal normal function. Dysfunctional mitochondria show impaired oxidative phosphorylation (OXPHOS), impaired mitochondrial dynamics and increased ROS production ^16^. Since brain does not have strong ability of antioxidant ^50^, increased oxidative damage can cause catastrophic damage to neurons ^13^. Mitochondrial swelling is accompanied by mitochondrial dysfunction during aging ^51^, and evidence suggests the contribution of pathological p-Tau on this process ^16^. Through pathological tau induced mitochondrial elongation in both *Drosophila* and mouse neurons ^18^, and EM images of P301S tauopathy mice showed tau aggregation in neurons with decreased mitochondria number ^6^, there is still lack of direct evidence to prove the role of p-Tau in mitochondrial swelling. In this study, by using electron microscopy analysis, we were able to observe swollen mitochondria in AT8-positive neurons in *APOE4* KI P301S tauopathy mice. Our data first showed mitochondrial swelling was increased in neurons with p-Tau aggregation, providing more direct evidence about the role of p-Tau on mitochondrial swelling. Through *in vitro* study using MSP-1 newborn neurons, we found that increased mitochondrial swelling would result in mitochondrial dysfunction, such as reduced neuronal mitochondrial membrane potential and increased oxidative response. Moreover, we found that mitochondrial dysfunction was also occurred in newborn neurons in E4-3Tg mice by RNA-Seq analysis of PSA-NCAM^+^ newborn neurons. Our data also showed decreased expression of genes related to neuron maturation represented by synapse plasticly, which was also found in the immature neurons of AD patients ^52^. AHN dysfunction in AD is associated with the impairment of maturation process ^47,53,54^, our findings suggested that mitochondrial abnormality and dysfunction may one of the reasons, which was a consequence of p-Tau. Taken together, our study revealed the contribution of p-Tau on neuronal mitochondrial swelling and dysfunction, both in mature neurons and newborn neurons.

Besides, our findings also showed that increased neuronal mitochondrial swelling led to neurodegeneration via activating necroptosis pathology. Impaired mitochondrial function and increased oxidative damage response (ROS) may account for this. Pathological tau is regarded as a main trigger for neuronal necroptosis ^21,24,38^, this study provided a possible explanation that tau-associated mitochondrial swelling and dysfunction may represent a key upstream step in this process. Noticeably, our findings showed the activation of necroptosis pathway in newborn neurons, which can be prevented by necroptosis inhibitor Nec-1. To date, this is the first report that newborn neurons would loss through the necroptosis pathway during the maturation process and provide a new insight of rescuing AHN levels by targeting mitochondrial function.

In addition, our study revealed that down-regulation of astrocytic C3 production can alleviate neuronal mitochondrial swelling and dysfunction in different tauopathy models via attenuating C3- C3aR signaling. Various studies have provided the evidence of the contribution of activated astrocytes on neuronal oxidative stress and neurodegeneration ^27^. Plasma GFAP levels has been regarded as a useful biomarker for dementia prediction ^55^. These suggest that astrocyte activation may play a crucial role in dementia progression. Complement system over-activation is accompanied by astrocyte activation with C3 over-production, which would promote tau pathology through C3-C3aR signaling via GSK3β pathway ^28–30^. We also observed that C3 expression had a close correlation with NFT number in TE4 mice and found the ability of C3 to increase p-Tau expression in MSP-1 newborn neurons. Therefore, down-regulation of astrocytic C3 attenuated tau phosphorylation, leading to alleviated mitochondrial swelling and dysfunction. Moreover, we reported the association of astrocytic C3 with neuronal necroptosis activation. Since blocking of astrocytic C3 production or attenuated C3- C3aR signaling have been reported to prevent tau-mediated neuronal loss ^30,32^, our findings suggested that astrocytic C3 may mediate neuronal mitochondrial swelling and dysfunction to cause necroptosis-related neuronal loss. Moreover, down-regulation of C3 production prevented the necroptosis of newborn neurons, and this explained the previous report that C3 affect the survival of newborn neurons^33^. To date, this study may be the first report of the impact of astrocytic C3 on neuronal mitochondrial swelling, dysfunction and subsequent necroptosis activation. Our findings filled the knowledge gap about the specific mechanisms between increased astrocytic C3 production and neuronal degeneration in tauopathy. In this study, we used a natural anti-inflammatory imidazole dipeptide (anserine) to block astrocytic C3 production. In previous study, we reported the ability of anserine on suppressing astrocyte activation ^39^, and here we further showed that anserine can target astrocytes by *Pept2* and suppress Tak1-IKK dependent NF-κB activation. Tak1 or Ikk activation in astrocytes occurs in many other neurodegeneration diseases ^56^, therefore, we believed that anserine may have more application value not only in AD and tauopathies and this need to be clarified in the future study.

Except for alleviating mitochondrial swelling, we found an interesting thing that down-regulate astrocytic C3 production prevented the cognitive decline and mortality both in TE4 and E4-3Tg mice. Reduced lifespan is also one of the characteristics of tau pathology due to the decreased muscle strength and paralysis ^6^. From our research, APP/PS1 model mice do not exhibit this phenotype, and we thought it caused by tau-related neuronal mitochondrial swelling and dysfunction. Research has indicated that p-Tau aggregation in the spinal cord leaded to the death of motor neurons, which may account for the decreased muscle strength ^57^. Reactive astrocytes have been observed in the spinal cord in tauopathy mice ^58^, therefore, suppressed astrocytic C3 may also alleviate the neuronal mitochondrial swelling in the spinal cord. Anyway, our findings revealed the harmful consequence of neuronal mitochondrial swelling, that is neurodegeneration, necroptosis activation and mortality because of mitochondrial dysfunction and increased oxidative stress. From a recent research, inhibition of neuronal necroptosis did not reduced mortality in tauopathy mice ^38^, while immunosuppressive agent tacrolimus (FK506) significantly prevented mortality ^6^. This suggested that preventing the mitochondrial swelling itself is more meaningful than preventing the consequences of mitochondrial swelling. In this study, we suppressed astrocyte activation and C3 production to alleviate tau-associated neuronal mitochondrial swelling and protected the neuronal function, and there may have some more direct approaches. Some studies have shown that mitochondrial swelling is due to the impairment Drp1 mediated mitochondrial fission, which would lead to dysfunction of mitophagy ^59–61^. Moreover, enhancing neuronal mitophagy can rescue mitochondrial dysfunction and attenuate tau pathology both in culture neurons and tauopathy model mice ^62,63^. Therefore, mitophagy maybe a potential target to alleviate the harmful effects of mitochondrial swelling.

In summary, our findings in this study revealed the impact of astrocytic C3 on mediating neuronal mitochondrial swelling, dysfunction and necroptosis activation and indicated that therapies that down- regulate astrocyte activation such as anserine have a neuroprotective role to alleviate this.

## Methods

### Animals

APPswe/PSEN1dE9 (B6/C3) and P301S tau (B6/C3) transgenic model mice were purchased from Jackson Laboratories (Bar Harbor, Maine, USA). APP/PS1 model mice express the Swedish variation of the phenotype, presenting both a chimeric human APP transgene (Mo/HuApp695swe) and a human PS1 transgene (missing exon 9). P301S tau transgenic model mice expressing human P301S 1N4R tau driven by PrP promoter. Human *APOE4* KI (C57BL/6) model mice were purchased from Riken BRC (Koyadai, Tsukuba, Japan). *APOE4* KI mice were crossed to APP/PS1 and P301S tau model mice to generate Tau or APP/PS1/Tau mice in human *APOE4* background, called TE4 and E4- 3Tg model mice. Only *APOE4* homozygote mice were used for experiment. *APOE4* KI mice were used as control. All the mice were sorted by age and genotype, kept under a 12-hour light / dark cycle at ∼22 °C, and could get food (solid feed MF; Oriental Yeast Co., Ltd., Tokyo, Japan) and water freely.

All animal procedures and experiments in this study were approved by the ethical committee of the University of Tokyo and were conducted according to the guidelines for animal experimentation required by the University of Tokyo.

### Anserine treatment

Anserine used in this experiment was purified from salmon muscles (purity>93%, Tokai Bussan, Tokyo, Japan), which only contains anserine and no carnosine. Anserine-treated mice were maintained on a steady dosage of anserine diluted in autoclaved drinking water, at a concentration of 2.0 g/L (10 mg/mouse per day) for 8 weeks. For TE4 mice, anserine treatment was performed from about 7 months of age to 9 months of age. For E4-3Tg mice, anserine treatment was performed from about 4 months of age to 6 months of age. Behavior tests were performed after the end of treatment.

### Y-maze

Y-maze was performed to assess short-term spatial memory of mice, which is suitable to show spontaneous alternation behavior. During the 10-min test, mice were placed on starting arm and allowed to explore the maze freely. The sequence and total number of arms entered were recorded by SMART video tracking system (Panlab, Barcelona, Spain). When mice enter 3 different arms in sequence, it is considered an alternation. The alternation rate was calculated as (the total number of arms entered minus 2 / maximum alternations) × 100. After each test session, the maze was cleaned with 70% ethanol solution.

### Radial arm maze

Radial arm maze (RAM) was performed to test spatial reference memory of mice. Mice were given limited food (2g /mouse per day) for at least three days before the test until the end of the test. In order to let the mice adapt the maze, mice were allowed to explore the maze and feeding freely for two days (habituation). There was a 20 mg reward in the end in all of the arms during the habituation. Then following the training trial. The training trial was performed for 5 days and two sessions daily with 1-hour intervals. Mice were placed on the start arm and were allowed to find all the rewards which were only placed in 3 arms. The time that mice find all the 3 rewards were recorded by SMART video tracking system (Panlab, Barcelona, Spain) or until 4 min had cost. After each test session, the maze was cleaned with 70% ethanol solution.

### Contextual fear conditioning test

Contextual fear conditioning (CFC) test was performed in P.O.BOX 319 (Med Associates Inc., Albans, UK) for 2 consecutive days. The design of the experiment was described previously ^64^. Conditioning test was assessed on the first day. Mice were placed into the box wiped with 70% isopropanol (Context A) for 370 seconds and given a sound (85 db, 5000 Hz, 10s) at 128 seconds, 212 seconds, and 296 seconds with an electric shock (0.75 mA, 1s) in the end. 24 hours after the Conditioning test, Context test was performed. Mice were placed into the same box (Context A) for 512 seconds and freezing time was recorded. The freezing behavior was recorded using Video Freeze Software (Med associate, Inc., St. Albans, UK).

### Grip strength test

Grip strength test was conducted to evaluate the physiological function of each mouse. Before the test, the grip strength metre (MK-380M; Muromachi Kikai) was cleaned with 70% ethanol solution. Mice were held by the tail and placed on the grip strength metre and make their two limbs grasped the metal grid. Then the operator gently pulled mice away in a direction horizontal to the grid until the two limbs were detached from the grid, and the peak pull-force was recorded. Each test was performed three sessions with 1 min intervals. The mean value was regarded as the grip strength of each mouse (two limbs). At the same time, the weight of each mouse was recorded to rule out the effect of body weight on grip strength.

### Immunohistochemical staining

Mice were perfused with TBS to remove blood and with 4% paraformaldehyde (PFA) to fix the brain tissue. Brain samples were taken and postfixed in 4% PFA for 24 hours, then incubated in 30% sucrose/TBS solution for 48 hours. Tissue-Tek OCT compound (Sakura Finetek, Japan) was used to embedded brain samples. All the brain samples were stored at −80 °C before use. Brain samples were sliced into 50-µm-thick coronal sections with a Cryostat (Microm, Germany) and kept in cryoprotectant solution at −30 °C. For immunofluorescence staining, sections were washed twice with 0.3% TBS-X (Triton) for 10 min and blocked with 3% normal donkey serum (NDS) diluted in 0.3% TBS-X for 60 min at room temperature and then incubated with primary antibodies overnight at 4°C with shaking. The primary antibodies used in this study were anti-GFAP (Mouse, 1:1000, Sigma), anti- C3 (Rat, 1:200, Hycult), anti-DCX (Rabbit, 1:500, Cell Signaling), anti-DCX (Sheep, 1:100, R&D Systems), anti-p-MLKL (Rabbit, 1:500, Abcam), anti-PT181 (Rat, 1:500, Wako), anti-AT8 (Mouse, 1:500, Thermo Fisher), anti-NeuN (Mouse, 1:500, Millipore), anti-BrdU (Rat, 1:100, Abcam) and anti-GFP (Rabbit, 1:500, MBL life science). Sections were then washed 3 times with TBS for 10 min and incubated with second antibodies for 2 hours at room temperature. The second antibodies used in this study were donkey anti-mouse IgG Alexa 647 (1:1000, Invitrogen), donkey anti-mouse IgG donkey Alexa 488 (1:1000, Invitrogen), goat anti-rabbit IgG Alexa 488 (1:1000, Invitrogen), donkey anti-rabbit IgG donkey Alexa 568 (1:1000, Invitrogen), goat anti-Rat IgG Alexa 594 (1:1000, Invitrogen) and donkey anti-sheep IgG NL557 (1:200, Jackson Immunoresearch). After wishing with TBS for 10 min, sections were incubated with DAPI (1: 10000, Sigma) in TBS for 5 min and then wished with TBS 2 times for 15 min. For NeuN staining, sections do not need to incubate with DAPI. Sections were finally mounted on microscope slides and visualized using a confocal microscope (FV3000-L4EN-TN21; Olympus, Japan). Specific information of antibodies used in this study can be seen in Supplementary Table. 1.

### Images analysis of immunohistochemical staining

Microscopy images were analyzed with ImageJ software. For stain area analysis like PT181 staining, immunoreactivity was quantified by the coverage area of the specific organization. The data was presented as the area % immunoreactivity ± S.E.M. per group. For cell counting analysis like p- MLKL and GFAP staining, the data was presented as the number of positive cells ± S.E.M. per group. For quantification of staining density in specific cells like C3, the staining area was calculated separately using ImageJ and then the staining density of the target factor in the cell was calculated as area %. More than 12 random cells were selected from 4 random acquisition areas for analysis of each mouse. The data was presented as the area in cells ± S.E.M. per group.

For each AT8 or PT181 staining image, AT8 or PT181 positive cells were counted as NFT. Data was presented as number of NFTs per section ± S.E.M. per group.

For counting the number of DCX-positive and BrdU-positive cells, the entire DG area was analyzed by moving the entire z-axis, and all images were collected. Each image corresponded to a thickness of 1.0 µm. All the images were opened used ImageJ, and then the image of the entire space was presented through the “stack-image to stack” step and “stack-Z project” step. The cell count average of each mouse was converted to the total number of cells in DG based on the standard length of the DG. To evaluate the maturation of newborn neurons, we defined mature DCX cells as DCX positive cells whose dendrites extend to the outside the granular cell layer (GCL). For p-MLKL and DCX co-staining, the data was presented as the number of co-labeling cells per section.

For all immunohistochemical staining, at least three random acquisition areas were considered for each brain section and more than two brain sections from Bregma −1.70 to 2.70 were analyzed for each mouse.

### Quantification of neuron density

Three brain sections (bregma −1.4, −1.7, and −2.0 mm) from each mouse were used to quantify the neuron density in CA1 and brain section (bregma −2.0 mm) form each mouse was used to quantify the neuron density in DG to exclude the influence of different positions of hippocampus. NeuN staining was performed to identify mature neurons. The numbers of NeuN positive cells were counted. Neuron density was presented as the number of NeuN positive cells in 1 mm^2^ ± S.E.M. per group.

### Sample preparation for Scanning Electron Microscopy

For observation of scanning electron microscopy (SEM), mice were perfused with TBS and fixed by 2% paraformaldehyde and 2% glutaraldehyde buffered with 0.1 M phosphate buffer (PB), (pH 7.2) as we previous described. Fixed brains were cut into 0.5-1 mm sections and hippocampus parts were obtained. Sections were then treated with 2% OsO4 in PB for 2h, dehydrated with a graded series of ethanol and embedded in epoxy resin (EPOK812, Okenshoji, Tokyo, Japan). Sections were cut into 90 nm thickness with the ultramicrotome (Leica UC7) and then stained with uranyl acetate and lead citrate. After coating, the sections were observed by electron microscope (Regulus 8240, Hitachi High-Tach Corporation, Tokyo, Japan). For the cell samples, fixed cells were treated with 1% OsO4 in PB for 2h and 0.5% uranyl acetate for 30min. Then the samples were dehydrated with a graded series of ethanol and embedded in epoxy resin, cut into 300 nm thickness with the ultramicrotome and stained for SEM observation as we describe above.

### Preparation for in-resin correlative light and electron microscopy

For in-resin correlative light and electron microscopy (CLEM), mice were perfused with TBS and fixed by 4% paraformaldehyde and 0.25% glutaraldehyde buffered with 0.1 M PB. After post-fixation overnight with the same solution, brains were stored in 0.1 M PB at 4°C before use. The brains were then sliced into 50-µm-thick sections by semiauto-matic Leica VT1200 vibrating blade microtome and stored in 0.05% Azide buffered with 0.1 M PB at 4°C. Brain sections were stained with primary antibodies and second antibodies and followed by post-fixation by 2% paraformaldehyde and 2% glutaraldehyde buffered with 0.1 M PB for at least 1h. Then brain sections were treated with 1% OsO4 in PB for 15 min, dehydrated with a graded series of ethanol and embedded in epoxy resin (EPOK812, Okenshoji, Tokyo, Japan). Sections were cut into 300 nm thickness with the ultramicrotome (Leica UC7) and observed by Nikon A1RHD25 confocal laser-scanning microscope to identify the target cells. Finally, the same sections were stained with uranyl acetate and lead citrate and observed by electron microscope (Regulus 8240, Hitachi High-Tach Corporation, Tokyo, Japan).

### Astrocytes and newborn neurons isolation

Magnetic associated cell sorting (MACS) was used to isolate astrocytes and newborn neurons from mouse brain. Mice were sacrificed under deep anesthesia and brains were removed carefully and washed in cold DPBS to remove the blood. Then cortex and hippocampus parts were taken and dissociated into single cell suspension by using Adult Brain Dissociation Kit (Miltenyi Biotec) based on the manufacturer’s protocol. For newborn neurons isolation, only hippocampus part was used. Then the astrocytes or newborn neurons were isolated using Anti-ACSA-2 MicroBead Kit or Anti-PSA-NCAM MicroBead Kit (Miltenyi Biotec). Briefly to say, the single cell suspension was incubated with mouse Fcg receptor block reagent at 4°C for 10 min following by anti-ACSA-2 or anti-PSA-NCAM MicroBead at 4°C for 15 min. Targeted cells were collected by MACS with an MS column (Miltenyi Biotec).

### Adeno-associated virus preparation

The packaging and purification of AAV-shPEPT2 and AAV-Scramble RNA were performed using TransIT-VirusGEN® Transfection Reagent (Mirus) and AAVpro® Purification Kit (Takara Bio, Japan) based on the manufacturer’s protocol. pAAV-U6-shPEPT2-EGFP vector and pAAV-U6- scramble_RNA vector were bought from VectorBulider, pHelper vector was a kind gift from Dr. Haruo Okado (Tokyo Metropolitan Institute of Medical Science), and pUCmini-iCAP-PHP.eB vector was bought from addgene (USA). AAV packaging was conducted in HEK293T cells.

### Intracerebroventricular injection

Intracerebroventricular injection was performed at about 6.5 months of age. TE4 mice were deep anesthesia with a mixed narcotic solution (0.3 mg/kg medetomidine, 4 mg/kg midazolam, and 5 mg/kg butorphanol tartrate, i.p.). Then mice were received injection of 4ul AAV-PHP.eB-U6-ShPEPT2-EGFP or 4ul AAV-PHP.eB-U6-Scramble_RNA-EGFP at both sides of ventricle using following coordinates: anterior-posterior (AP) = −0.4 mm; medial-lateral (ML) = ±1.0 mm; dorsal-ventral (DV) = −2.0 mm). The injection was performed using an auto-nanoliter injector (Nanoject II, Drummond SCI, USA), and the needle was kept at least 5 min after the injection to prevent the backflow of AAV solution. Each mouse was allowed to recover for 2 weeks and anserine treatment would begin at 7 months of age as described above.

### Nec-1 treatment

For nec-1 treatment in E4-3Tg mice, nec-1 (Selleck) was dissolved in 5%DMSO, 45%PEG300 and 50% ddH_2_O solution. Each mouse was received intraperitoneal injection with the dose of 6.25mg/Kg as previous described twice a week for 8 weeks ^65^. After the last injection, the behavior tests were performed.

### Mouse striatal precursor-1 (MSP-1) cells culture

MSP-1 cells line was obtained from a neural stem cell line established from ventral telencephalon tissue of p53 KO mice as we previous described ^42^. MSP-1 cells were cultured in poly-L-lysine/fibronectin-coated (Sigma) dishes with DMEM/F12 medium (Gibco) containing 10% FBS (Gibco) and fibroblast growth factor (basic-FGF) (10 ng/ml, Wako). This medium was called MSP-1 proliferation medium. Culture medium was replaced with fresh medium every 2 or 3days.

To obtain differentiated astrocytes, MSP-1 cells were harvested and replaced on new dishes overnight with MSP-1 proliferation medium. The next day, replaced the medium with serum-free DMEM/F12 medium containing N-2 supplement (R&D system), Leukemia Inhibitory Factor (LIF) (10 ng/ml, Wako) and Ciliary Neurotrophic Factor (CNTF) (10 ng/ml, Wako). This medium was called astrocytes-differentiation medium. Culture medium was replaced with fresh medium every 2 or 3 days. On day 5, cells were exposed to Tumor Necrosis Factor-α (TNF-α) (100ng/ml, Wako) for 24 hours. For anserine group, cells were pre-treated with L-Anserine (Toronto Research Chemicals) for 2 hours before exposed to TNF-α.

To obtain differentiated neurons, MSP-1 cells were replaced the medium with serum-free DMEM/F12 medium containing N-2 supplement. This medium was called neurons-differentiation medium. Culture medium was replaced with fresh medium every 2 days. On day 5, cells were treated with cultured supernatant of MSP-1 differentiation-induced astrocytes or mouse C3/C3a protein for 24 hours. Nec-1 and C3aR antagonist (SB290157) were bought from Selleck. Mouse C3/C3a protein was bought from MedChemExpress. Regents’ information can be seen in Supplementary Table. 2.

### Proteins isolation

For isolation of brain proteins, mice were perfused by TBS and hippocampus parts were taken. Tissues were lysed by RIPA buffer containing 1 mM PMSF (Cell signaling), Protease Inhibitor Cocktail (Wako) and Phosphatase Inhibitor Cocktail (Wako) on ice for 1-2h. For cell samples, harvested cells were lysed by the same RIPA buffer on ice for 30 min. Then, the lysates were centrifuged at 15,000g for 30 min at 4°C and the supernatants were collected (RIPA-soluble fraction). Protein samples were stored at -80 °C before use.

### Western-Blot

Equal isolated proteins (20-40ug) were electrophoresed by sodium dodecyl sulfate-polyacrylamide gel electrophoresis (SDS-PAGE) and transferred proteins from gel onto polyvinylidene difluoride fluoride (PVDF) membranes (BioRad). After transformation, the blotting membranes were blocked with EveryBlot blocking buffer (BioRad) with shaking for 5 minutes at room temperature. Subsequently, blocked membranes were incubated with diluted primary antibodies at 4 °C overnight. The primary antibodies used in this study were anti-*α*-Tubulin (Mouse, 1:2000, Proteintech), anti-Tak1 (Rabbit, 1:500, Cell signaling), anti-p-Tak1 (Rabbit, 1:500, Cell signaling), anti-IKK (Rabbit, 1:1000, Proteintech) and anti-p-IKK (Rabbit, 1:500, Cell signaling). All the membranes would be washed with 0.05% TBS-T for five cycles for five minutes each, the membranes were then incubated with diluted secondary antibodies for one hour at 37 ℃. The secondary antibodies used in this study were HRP-conjugated Goat Anti-Mouse IgG (1:10000, Proteintech) and HRP-conjugated Goat Anti-Rabbit IgG (1:10000, Proteintech). Following, the membranes processed the same washing procedure as before. Finally, Clarity Max Western ECL Substrate (BioRad) and Image Gauge version 3.41 (Fuji Film, Japan) was used to detect the signals. Relative expression of target protein was analyzed by Image J using *α*-Tubulin as a host protein. Specific information of antibodies used in this study can be seen in Supplementary Table. 1.

### Immunocytochemistry staining

Harvested MSP-1 cells were cultured in poly-L-lysine/fibronectin-coated chamber glass (4 wells or 8 wells, Watson) with MSP-1 differentiation medium to get astrocytes or neurons differentiation. After treatment, cells were fixed with 4% PFA for 10 min and following a wash in TBS for 10 min. Then cells were blocked with 5% BSA diluted in 0.05% TBS-T for 30 min at room temperature and then incubated with blocking solution contained primary antibodies overnight at 4°C. The primary antibodies used in this study were anti-AT8 (mouse, 1:1000, Thermo fisher), anti-PSA-NCAM (mouse, 1:1000, gift from Dr. Seki), anti-MAP2 (Rabbit, 1:1000, Proteintech), anti-p-p65 (rabbit, 1:500, Cell signaling), anti-p-Ikk (Rabbit, 1:1000, Cell signaling), anti-p-Tak1 (rabbit, 1:1000, Cusabio) and anti-p-MLKL (Rabbit, 1:2000, Abcam). Cells were then washed 3 times with TBS for 5 min and incubated with second antibodies for 2 hours at room temperature. The second antibodies used in this study were donkey anti-mouse IgG Alexa 488 (1:1000, Molecular Probes), goat anti-mouse IgM Alexa 488 (1:1000, Abcam) and donkey anti-rabbit IgG Alexa 568 (1:1000, Molecular Probes). After wishing with TBS for 5 min, sections were incubated with DAPI (1: 10000, Sigma) in TBS for 5 min and then wished with TBS 2 times for 5 min. Cells were visualized using a confocal microscope (FV3000- L4EN-TN21; Olympus, Japan). Specific information of antibodies used in this study can be seen in Supplementary Table. 1.

### Image analysis of immunocytochemistry staining

Microscopy images were analyzed with ImageJ software. For AT8, p-MLKL, p-Tak1 and p-IKK staining, the stain intensity was calculated. The data was presented as the relative intensity (radio to sham) ± S.E.M. per group. For p-p65 staining, only the stain intensity in nucleus was calculated and the data was presented as the relative intensity ± S.E.M. per group. More than 100 cells were analyzed in each group from at least 10 random images.

### Mitochondrial function assay

Mitochondrial membrane potential, mitochondrial morphology and oxidative stress in MSP-1 neurons were analyzed using MT-1 MitoMP Detection Kit (Dojindo), Mitobright IM Red (Dojindo) and ROS Assay Kit -Photo-oxidation Resistant DCFH-DA- (Dojindo) according to the manual. Briefly to say, after the CM treatment, cells were treated with corresponding reagents for 30 min and then washed with HBSS. After fixed with 4% paraformaldehyde, Cells were visualized using a confocal microscope (FV3000-L4EN-TN21; Olympus, Japan). Obtained imaged were analyzed by Image J.

### RNA sequencing

Total RNA from isolated astrocytes or newborn neurons and MSP-1 differentiation induced astrocytes was isolated using RNeasy Mini Kit (Qiagen).1ng (isolated astrocytes and newborn neurons) or 10ng (cultured astrocytes) RNA samples were pre-treated with SMART-seq stranded Kit (Takara Bio, Japan). The sequencing was performed using Illumina NovaSeq6000 (Illumina, USA).

Transcripts per million (TPM) data was used to evaluate the genes expression levels and different expressed genes (DEGs) were identified by DESeq2 package in R environment. DEGs were further used for Gene Ontology (GO) analysis and Kyoto Encyclopedia of Genes and Genomes (KEGG) pathway analysis using David analysis (https://david.ncifcrf.gov/). Gene set enrichment analysis (GSEA) was performed based on the TPM data using GESA software (UC San Diego and Broad Institute). The figures were generated by ggplot2 package in R environment.

### Real-time quantitative PCR (RT-qPCR)

Isolated newborn neurons and astrocytes RNA samples (1-2ng) or cultured MSP-1 astrocytes samples (500ng) were used for Reverse transcription reaction for cDNA synthesis using SuperScript™ III Reverse Transcriptase (Thermo Fisher) according to the instruction manuals. RT-qPCR was conducted using TP-850 (Takara Bio, Japan) and TB green Taq (Takara Bio, Japan) with following protocol: an initial 30s denaturation at 95°C, then 5s at 95°C and 30s at 60°C for 50 cycles because of the low concentration of the samples. All the samples were run into double and β-actin was used as a reference gene. Analysis was performed with the 2^−ΔΔCt^ method and expressed as fold changes. The gene sequences for RT-qPCR were designed Primer 3 (Supplementary Table. 3).

## Statistical analyses

All Data were expressed as the mean ± SEM and were analyzed with GraphPad Prism (version 8; GraphPad Software). A two-tailed Student’s t-test was used for the statistical comparison of two samples. For groups more than 3, a one-way ANOVA with Tukey post-hoc test was used. For data across days, data were analyzed by two-way ANOVA with a Holm-Sidak post-hoc test was used for Multiple comparison test. A p value less than 0.05 was considered statistically significant.

## Acknowledgements

We thanked Tokai Bussan for the anserine sample. We thanked Mrs. Izumi Shiraishi and Mrs. Keiko Murasakino for the daily support in the lab. We thanked the gift of pHelper vector from Dr. Haruo Okado. We thanked Dr. Tatsunori Seki for the kindly gift of PSA-NCAM antibody. In additional, this work was supported by JST SPRING, Grant Number JPMJSP2108. Some of the diagrams were created with BioRender.com.

## Author Contributions

C.L., I.T., Y.U, K.S. and T.H. designed the experiments. C.L., B.Z., J.Y., R.T., Y.S. and M.S. performed the experiment and collected the data. Y.L., X.C., J.Y., I.T., Y.U and K.S. prepared the reagents, materials and instruments. C.L. and B.Z. analyzed the data and prepared the figures. C.L., B.Z., I.T., Y.U and T.H prepared the manuscript with the assistance of other authors. All authors have read this manuscript and approved the manuscript.

