## supplementary figure for "Impact of astrocytic C3-production on neuronal mitochondrial dysfunction in tauopathy mouse models"

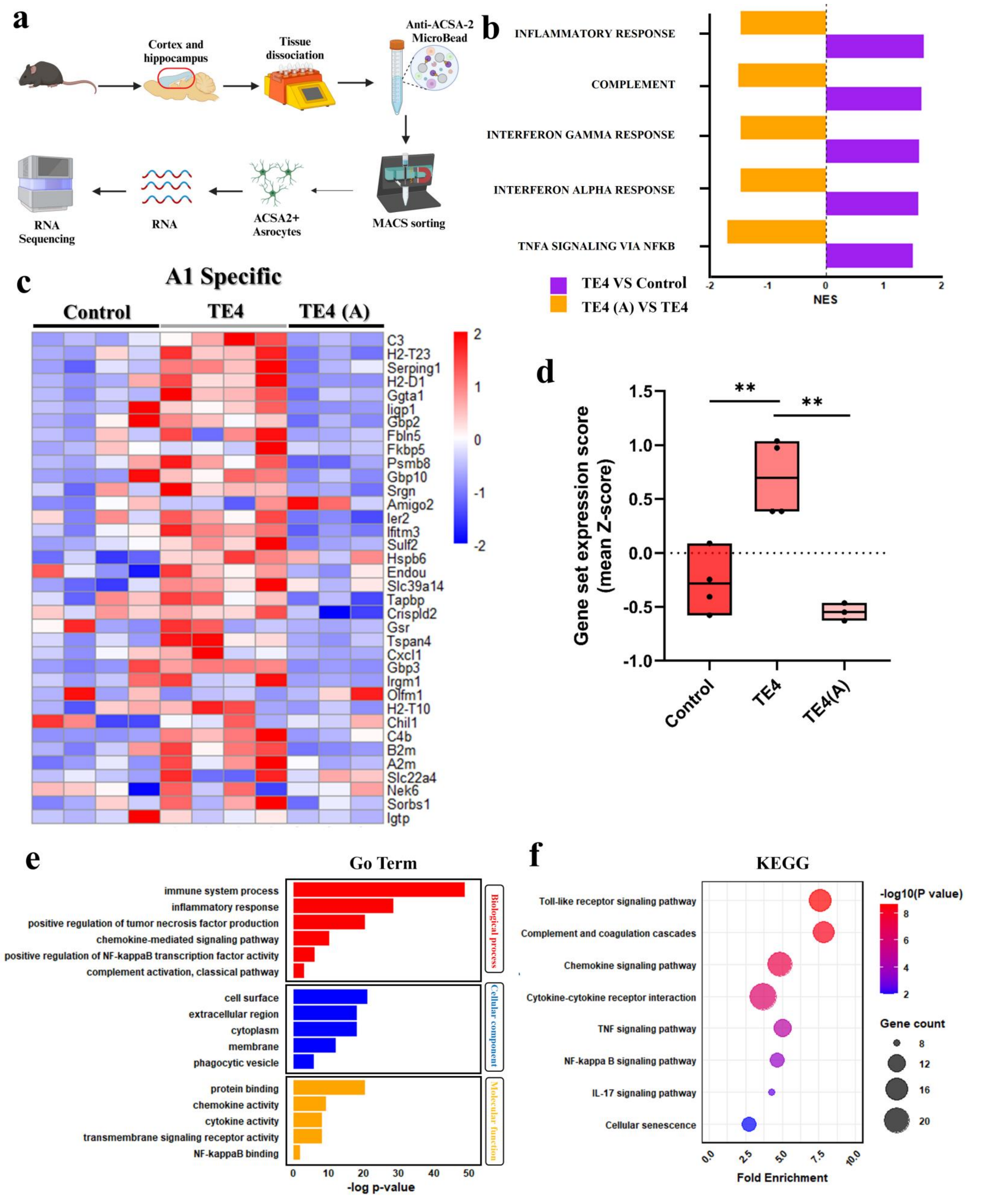


**Supplementary Fig. 1: Down-regulation of A1-phenotype astrocyte activation by a natural anti-inflammatory peptide, anserine, in TE4 mice.**

**a** The description of astrocytes isolation for RNA-Seq used ACSA-2 Microbead from mouse brain. **b** GESA results using the Hallmark database. The top 5 up-regulated gene sets were selected. **c** Heatmaps that exhibit expression levels of selected A1 specific astrocyte markers in control, TE4 and TE4 (A) mice. **d** Quantification of gene set expression score of A1 specific astrocytes markers in each group (*n*=4). **e** and **f** Results of KEGG (**e**) and Go term analysis (**f**) using common DEGs. Data represent means ± SEM and were analyzed by one-way ANOVA with Tukey's multiple comparisons test. $^{**}p < 0.01$. Some elements of figure (**a**) were created in https://BioRender.com.

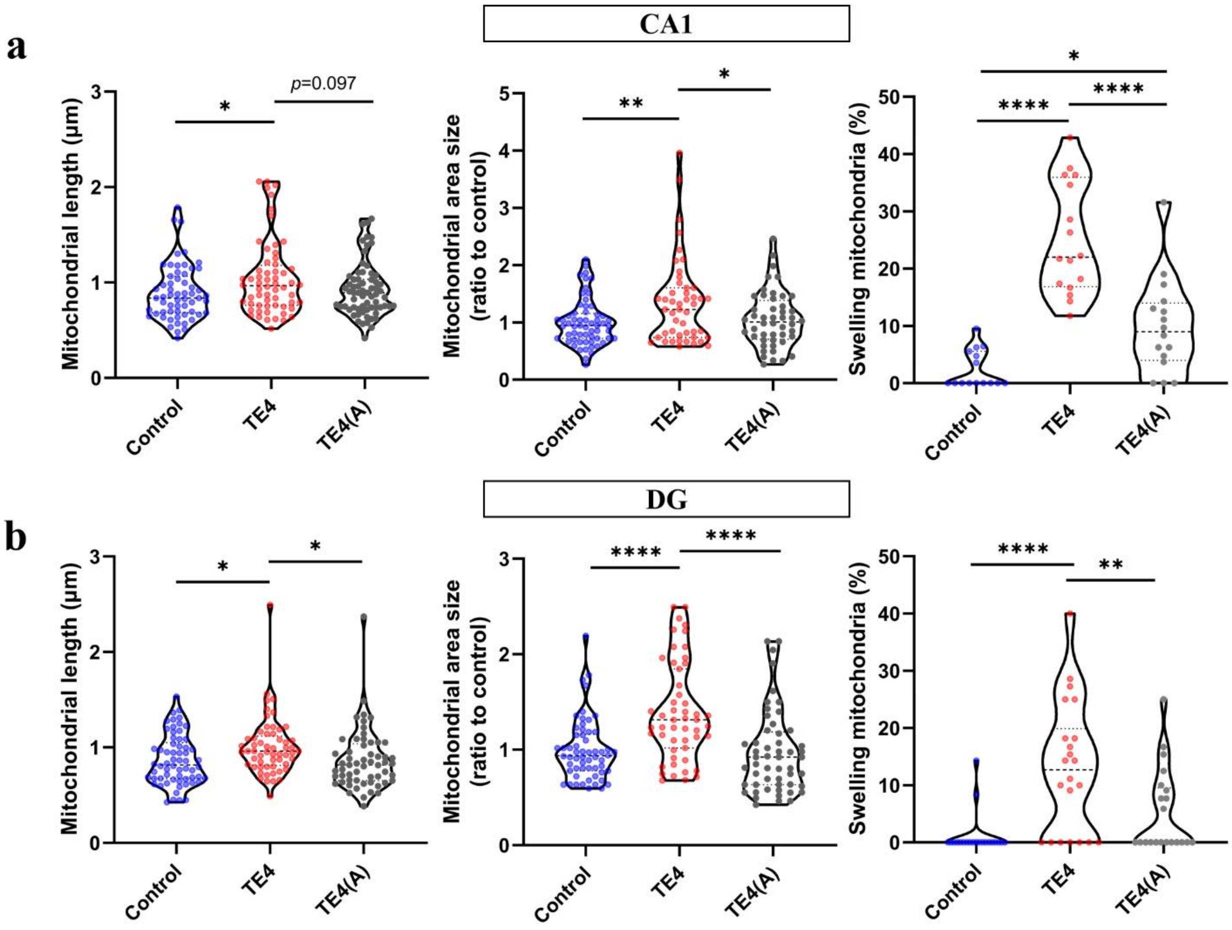


**Supplementary Fig. 2: Impact of astrocytic C3 production on neuronal mitochondrial swelling in TE4 mice.**

**a** Quantification mitochondrial length, mitochondrial area size and swelling mitochondria ratio in CA1 area of 3 groups of mice (neuron number: 15-16, mitochondria number: 47-67). **b** Quantification mitochondrial length, mitochondrial area size and swelling mitochondria ratio in DG area of 3 groups of mice (neuron number: 21-22, mitochondria number: 51-62). Data was showed in the violin plots (median: bold dashed line; quarters (1/4 and 3/4: dashed lines) and were analyzed by one-way ANOVA with Tukey's multiple comparisons test. *$p < 0.05$, **$p < 0.01$, ****$p < 0.0001$.

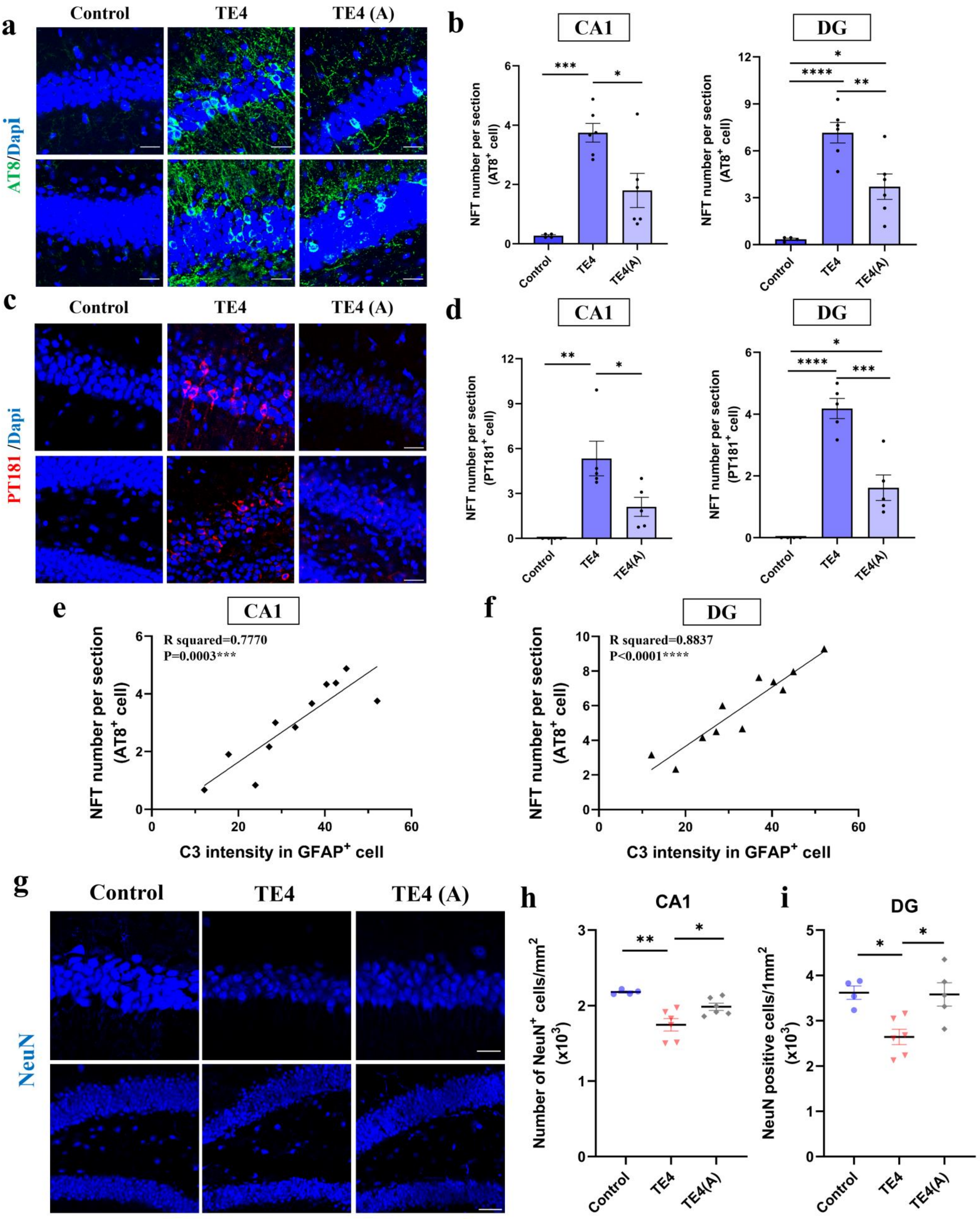
a
Control
TE4
TE4 (A)
AT8/Dapi
b
CA1
DG
NFT number per section (AT8+ cell)
Control
TE4
TE4(A)
c
PT181 /Dapi
d
NFT number per section (PT181+ cell)
e
CA1
R squared=0.7770
P=0.0003***
NFT number per section (AT8+ cell)
C3 intensity in GFAP+ cell
f
DG
R squared=0.8837
P<0.0001****
g
NeuN
h
CA1
Number of NeuN+ cells/mm2 (x10³)
i
DG
NeuN positive cells/1mm2 (x10³)

**Supplementary Fig. 3: Impact of astrocytic C3 production on tau pathology and neuronal loss in TE4 mice.**

**a** Representative images of AT8 staining in CA1 and DG area of 3 groups of mice (scale bar: 25 μm). **b** Quantification of NFT number ($AT8^+$) CA1 and DG area in each group, (*n*=4-6). **c** Representative images of PT181 staining in CA1 and DG area of 3 groups of mice (scale bar: 25 μm). **d** Quantification of NFT number ($PT181^+$) in CA1 and DG area in each group, (*n*=4-5). **e** and **f** The simple linear regression curve of NFT number in CA1 (**e**) and DG (**f**) area and C3 intensity in GFAP-positive cells of TE4 mice, (*n*=11). **g** Representative images of NeuN staining in CA1 and DG area of 3 groups of mice (scale bar: 25μm (up), 50μm (down)). **h** and **i** Quantification of neuron density in CA1 (**h**) and DG area (**i**) in each group, (*n*=4-6). Data represent means ± SEM and were analyzed by one-way ANOVA with Tukey's multiple comparisons test. $*p < 0.05$, $**p < 0.01$, $***p < 0.001$, $****p < 0.0001$.

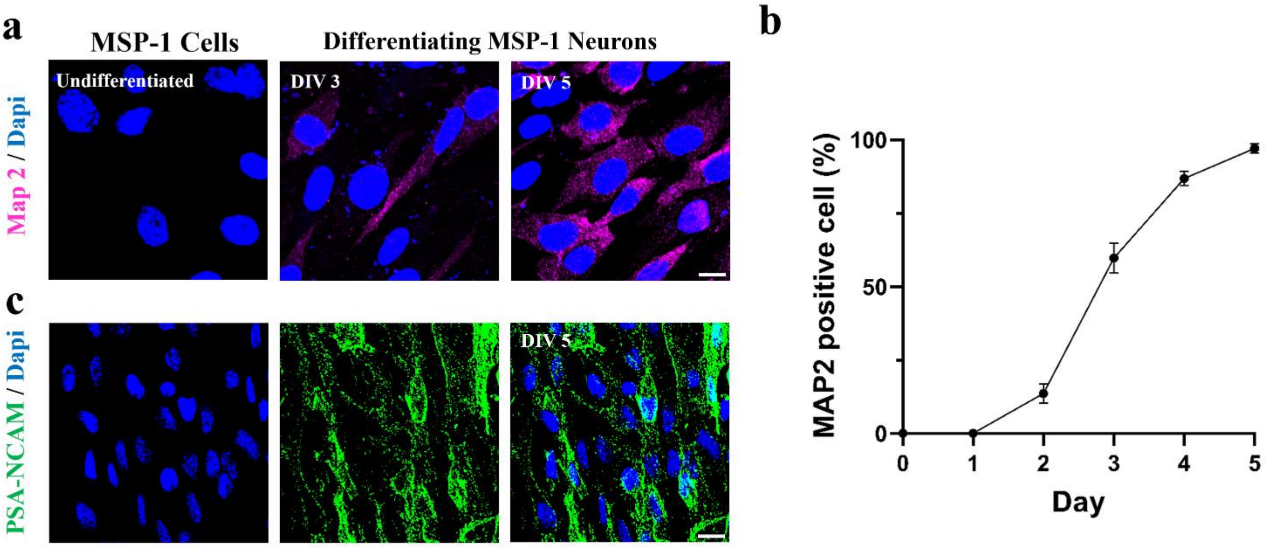


**Supplementary Fig. 4: Differentiation results of MSP-1 newborn neurons.**

**a** Representative images of MAP2 staining during the neuron-differentiation of MSP-1 cells (scale bar: 10 μm). **b** Quantification of MAP2-positive cells ratio during DIV0-DIV5, (*n*=3). **c** Representative images of PSA-NCAM staining in MSP-1 neurons at DIV5 (scale bar: 25 μm). Data represent means ± SEM.

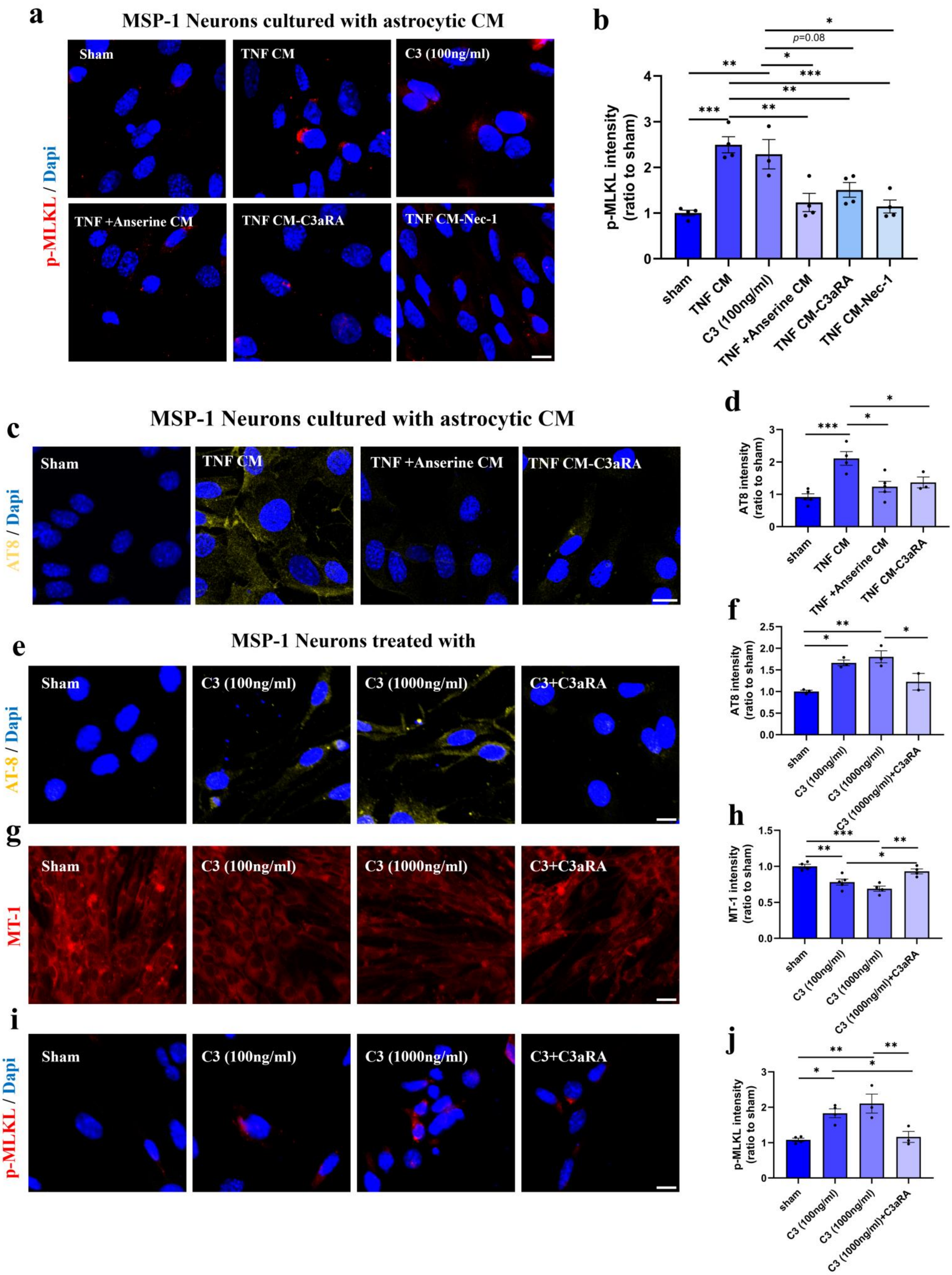

a
MSP-1 Neurons cultured with astrocytic CM
p-MLKL / Dapi
Sham
TNF CM
C3 (100ng/ml)
TNF +Anserine CM
TNF CM-C3aRA
TNF CM-Nec-1
b
p-MLKL intensity (ratio to sham)
p=0.08
sham
TNF CM
C3 (100ng/ml)
TNF +Anserine CM
TNF CM-C3aRA
TNF CM-Nec-1
c
MSP-1 Neurons cultured with astrocytic CM
AT8 / Dapi
Sham
TNF CM
TNF +Anserine CM
TNF CM-C3aRA
d
AT8 intensity (ratio to sham)
sham
TNF CM
TNF +Anserine CM
TNF CM-C3aRA
e
MSP-1 Neurons treated with
AT-8 / Dapi
Sham
C3 (100ng/ml)
C3 (1000ng/ml)
C3+C3aRA
f
AT8 intensity (ratio to sham)
sham
C3 (100ng/ml)
C3 (1000ng/ml)
C3 (1000ng/ml)+C3aRA
g
MT-1
Sham
C3 (100ng/ml)
C3 (1000ng/ml)
C3+C3aRA
h
MT-1 intensity (ratio to sham)
sham
C3 (100ng/ml)
C3 (1000ng/ml)
C3 (1000ng/ml)+C3aRA
i
p-MLKL / Dapi
Sham
C3 (100ng/ml)
C3 (1000ng/ml)
C3+C3aRA
j
p-MLKL intensity (ratio to sham)
sham
C3 (100ng/ml)
C3 (1000ng/ml)
C3 (1000ng/ml)+C3aRA

**Supplementary Fig. 5: Complement C3 induced tau-phosphorylation, mitochondrial dysfunction, and necroptosis activation in MSP-1 newborn neurons.**

**a** Representative images of p-MLKL staining of MSP-1 neurons treated with sham, TNF CM, C3(100ng/ml), TNF +Anserine CM, TNF CM-C3aRA (10μM) and TNF CM-Nec-1 (scale bar: 10 μm). **b** Quantification of p-MLKL intensity in each group, (*n*=3-4). **c** Representative images of AT8 staining of MSP-1 neurons treated with sham, TNF CM, TNF +Anserine CM and TNF CM-C3aRA (10μM) (scale bar: 10 μm). **d** Quantification of AT8 staining in each group, (*n*=3-4). **e** Representative images of AT8 staining of MSP-1 neurons treated with sham, C3 (100ng/ml), C3 (1000ng/ml) and C3-C3aRA (10μM) (scale bar: 10 μm). **f** Quantification of AT8 intensity in each group, (*n*=3-4). **g** Representative images of MT-1 staining of MSP-1 neurons treated with sham, C3 (100ng/ml), C3 (1000ng/ml) and C3-C3aRA (10μM) (scale bar: 25 μm). **h** Quantification of MT-1 intensity in each group, (*n*=4-5). **i** Representative images of p-MLKL staining of MSP-1 neurons treated with sham, C3 (100ng/ml), C3 (1000ng/ml) and C3-C3aRA (10μM) (scale bar: 10 μm) **j** Quantification of p-MLKL intensity in each group, (*n*=3-4). Data represent means ± SEM and were analyzed by one-way ANOVA with Tukey's multiple comparisons test. *$p < 0.05$, **$p < 0.01$, ***$p < 0.001$, ****$p < 0.0001$.

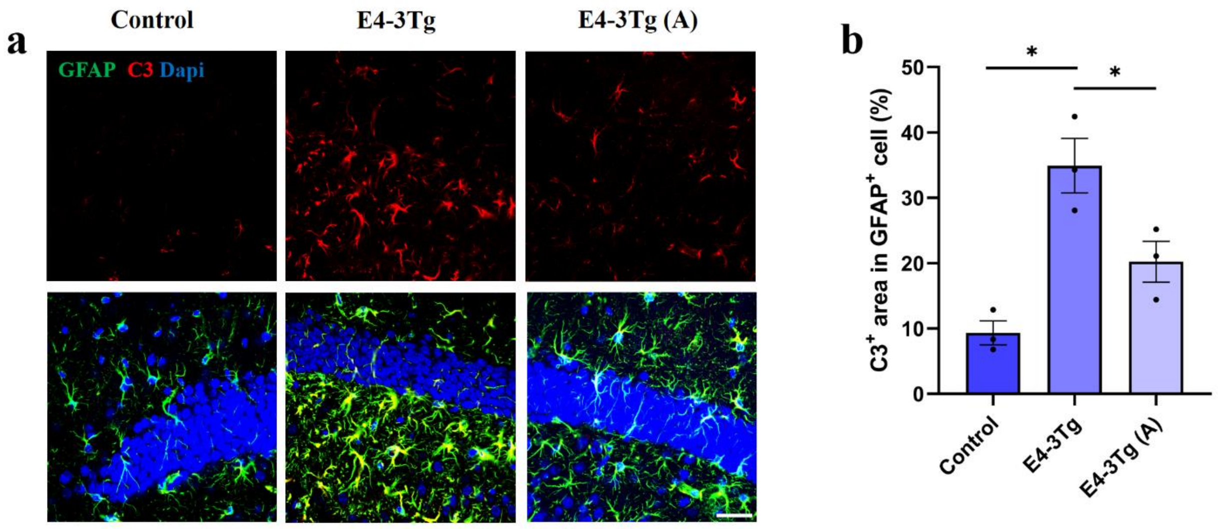


**Supplementary Fig. 6: Down-regulation of astrocytic C3 production in E4-3Tg (A) mice.**
**a** Representative images of GFAP and C3 co-staining in DG area of control, E4-3Tg and E4-3Tg (A) mice (scale bar: 25 μm). **b** Quantification of C3-positive area in GFAP-positive cells in each group, (*n*=3). Data represent means ± SEM and were analyzed by one-way ANOVA with Tukey's multiple comparisons test. *$p$<0.05.

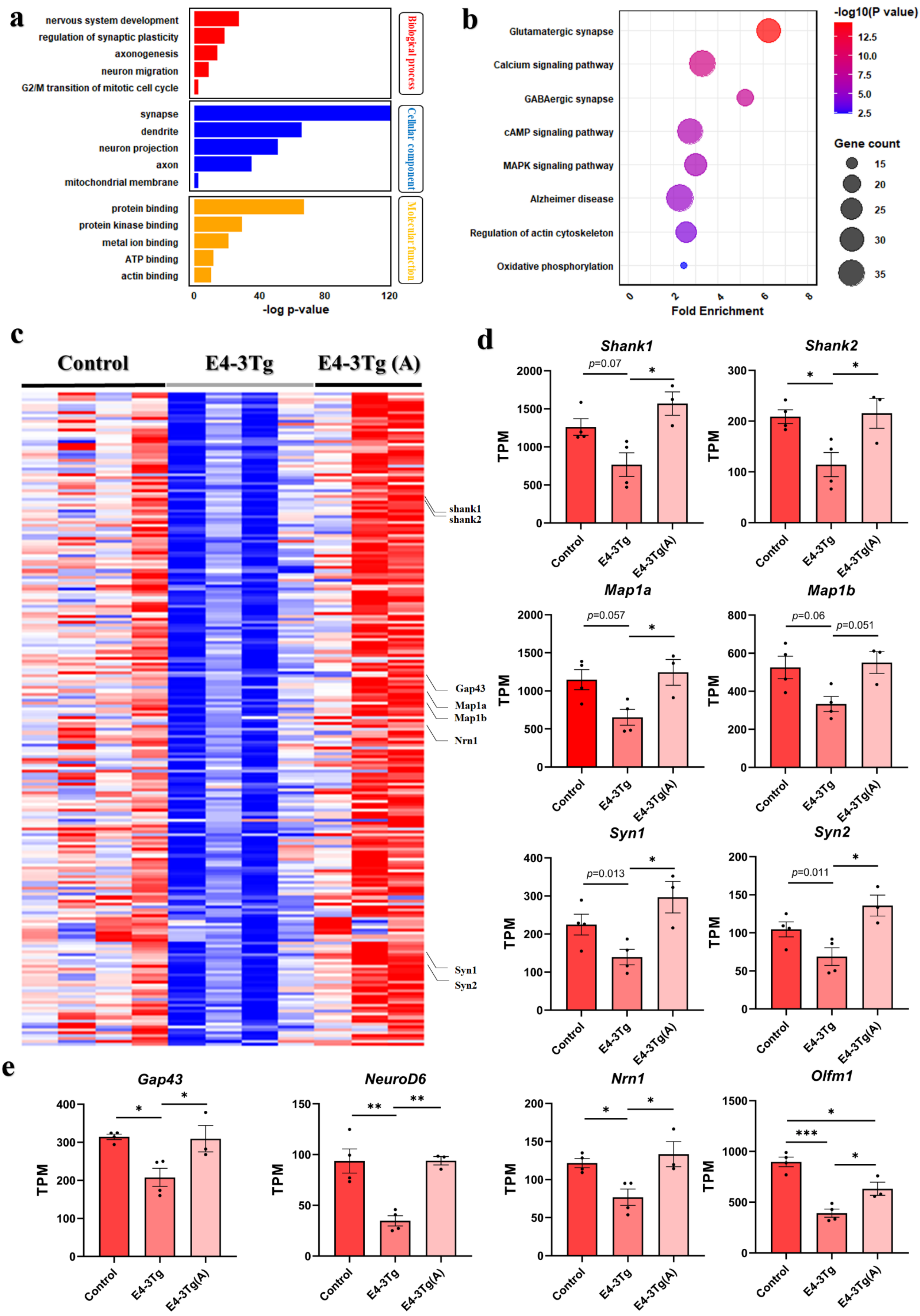

a
nervous system development
regulation of synaptic plasticity
axonogenesis
neuron migration
G2/M transition of mitotic cell cycle
synapse
dendrite
neuron projection
axon
mitochondrial membrane
protein binding
protein kinase binding
metal ion binding
ATP binding
actin binding
Biological process
Cellular component
Molecular function
-log p-value
b
Glutamatergic synapse
Calcium signaling pathway
GABAergic synapse
cAMP signaling pathway
MAPK signaling pathway
Alzheimer disease
Regulation of actin cytoskeleton
Oxidative phosphorylation
Fold Enrichment
-log10(P value)
Gene count
c
Control
E4-3Tg
E4-3Tg (A)
shank1
shank2
Gap43
Map1a
Map1b
Nrn1
Syn1
Syn2
d
Shank1
Shank2
Map1a
Map1b
Syn1
Syn2
p=0.07
p=0.057
p=0.06
p=0.051
p=0.013
p=0.011
TPM
Control
E4-3Tg
E4-3Tg(A)
e
Gap43
NeuroD6
Nrn1
Olfm1

**Supplementary Fig. 7: Dysfunction of Synaptogenesis and maturation process in newborn neurons of E4-3Tg mice.**

**a-b** Results of Go term (**a**) and KEGG analysis (**b**) using common DEGs that were down-regulated in E4-3Tg group compared with control group and up-regulated in E4-3Tg (A) mice compared with E4-3Tg mice. **c** Heatmap that shows expression levels of synapse markers of 3 groups of mice. All the selected genes were significantly down-regulated in E4-3Tg group and up-regulated in E4-3Tg (A) group. **d** and **e** Representative expression of genes related synaptogenesis (**d**) and neuronal maturation (**e**) in each group, ($n$=3-4). Data represent means ± SEM and were analyzed by one-way ANOVA with Tukey's multiple comparisons test. *$p < 0.05$, **$p < 0.01$, ***$p < 0.001$.

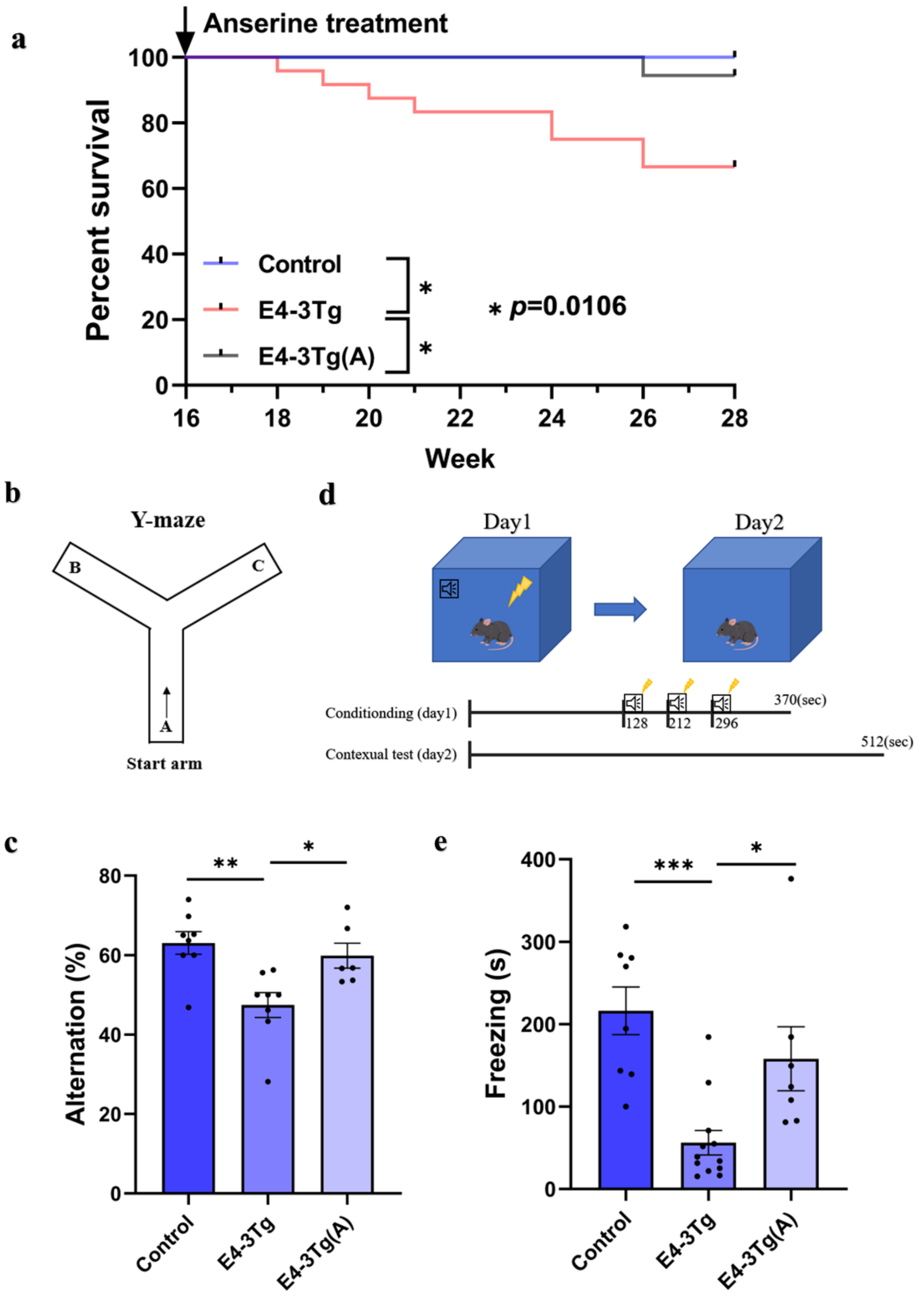


**Supplementary Fig. 8: Improved AHN rescued mortality and cognitive function in E4-3Tg mice.**
**a** Kaplan–Meier survival curve of control, TE4 and TE4 (A) groups, (*n*=12-24). Kaplan-Meier survival analysis was performed by log-rank (Mantel-Cox) test, $p$=0.0106. Control and E4-3Tg, $p$=0.0294; E4-3Tg and E4-3Tg (A), $p$=0.0289. **b** Description of Y-Maze, mice would start at A site. **c** Result of Y-maze. The result was presented as alternation rates (%) of each group of mice during 10 min, (*n*=6-8). **d** Description of CFC test. **e** Quantification of Freezing time in each group at contextual test, (*n*=7-12). Data represent means ± SEM and were analyzed by one-way ANOVA with Tukey's multiple comparisons test. *$p < 0.05$, **$p < 0.01$, ***$p < 0.001$.
